# AGO1x prevents dsRNA-induced interferon signaling to promote breast cancer cell proliferation

**DOI:** 10.1101/603506

**Authors:** Souvik Ghosh, Joao C Guimaraes, Manuela Lanzafame, Alexander Schmidt, Afzal Pasha Syed, Beatrice Dimitriades, Anastasiya Börsch, Shreemoyee Ghosh, Ana Luisa Correia, Johannes Danner, Gunter Meister, Luigi M. Terracciano, Salvatore Piscuoglio, Mihaela Zavolan

## Abstract

Initially reported for viral RNA, elongation of polypeptide chains beyond the stop codon (translational readthrough (TR)) also occurs on eukaryotic transcripts. TR diversifies the proteome and can modulate protein levels ^1-6^. Here we report that AGO1x, a conserved TR isoform of Argonaute 1, is generated in highly proliferative breast cancer cells, where it curbs accumulation of double stranded RNAs, the induction of the interferon response and apoptosis. In contrast to other mammalian Argonaute protein family members with primarily cytoplasmic functions, AGO1x localizes to the nucleus, in the vicinity of nucleoli. We identify a novel interaction of AGO1x with the Polyribonucleotide Nucleotidyltransferase 1, depletion of either protein leading to dsRNA accumulation and impaired cell proliferation. Our study thus uncovers a novel function of an Argonaute protein outside of the miRNA effector pathway, in buffering dsRNA-induced interferon responses. As AGO1x expression is tightly linked to breast cancer cell proliferation, our study suggests a new direction for limiting tumor growth.

## Introduction

Guided by small RNAs — miRNA or siRNA — the four members of the human Argonaute protein family repress translation and promote degradation of mRNA targets ^7^, with largely overlapping target specificities ^8^. Although Argonaute proteins are primarily found in the cytoplasm, associations with organelles, as well as their presence in the nucleus have also been reported. In particular, Argonaute 1 (AGO1 or EIF2C1) has been found at promoters and enhancers, modulating chromatin marks ^9^, transcription ^10,11^ and alternative splicing ^9,12^. *AGO1* has emerged among the best predicted substrates of heterogeneous nuclear ribonucleoprotein A2/B1-dependent, programmed translational readthrough (TR), which also generates a particular isoform of vascular endothelial growth factor A (VEGF-Ax) ^3,13^. In contrast to VEGF-Ax, the expression pattern and function of the AGO1 TR isoform (AGO1x) in human cells have not been studied.

## Results

Analyzing the conservation of genomic regions downstream of annotated stop codons among vertebrates, we found a few cases where highly conserved C-terminal protein extensions could be predicted (**Fig. 1a**). The proteins encoded by these transcripts have primarily RNA and protein binding potential and are involved in the regulation of gene expression and metabolism. Among Argonaute family members, AGO1 exhibits the strongest conservation downstream of the annotated stop codon (**Fig. 1a**). Further support for the coding potential of this region comes from the striking paucity of nonsense mutations in the 99 nts-long region extending to the next predicted stop codon, as well as a frame-preserving, 3-nt deletion in Tarsiers (**Fig. 1b**). The predicted peptide extension is also extremely conserved across vertebrates (**Fig. 1c**).

**Figure 1.**
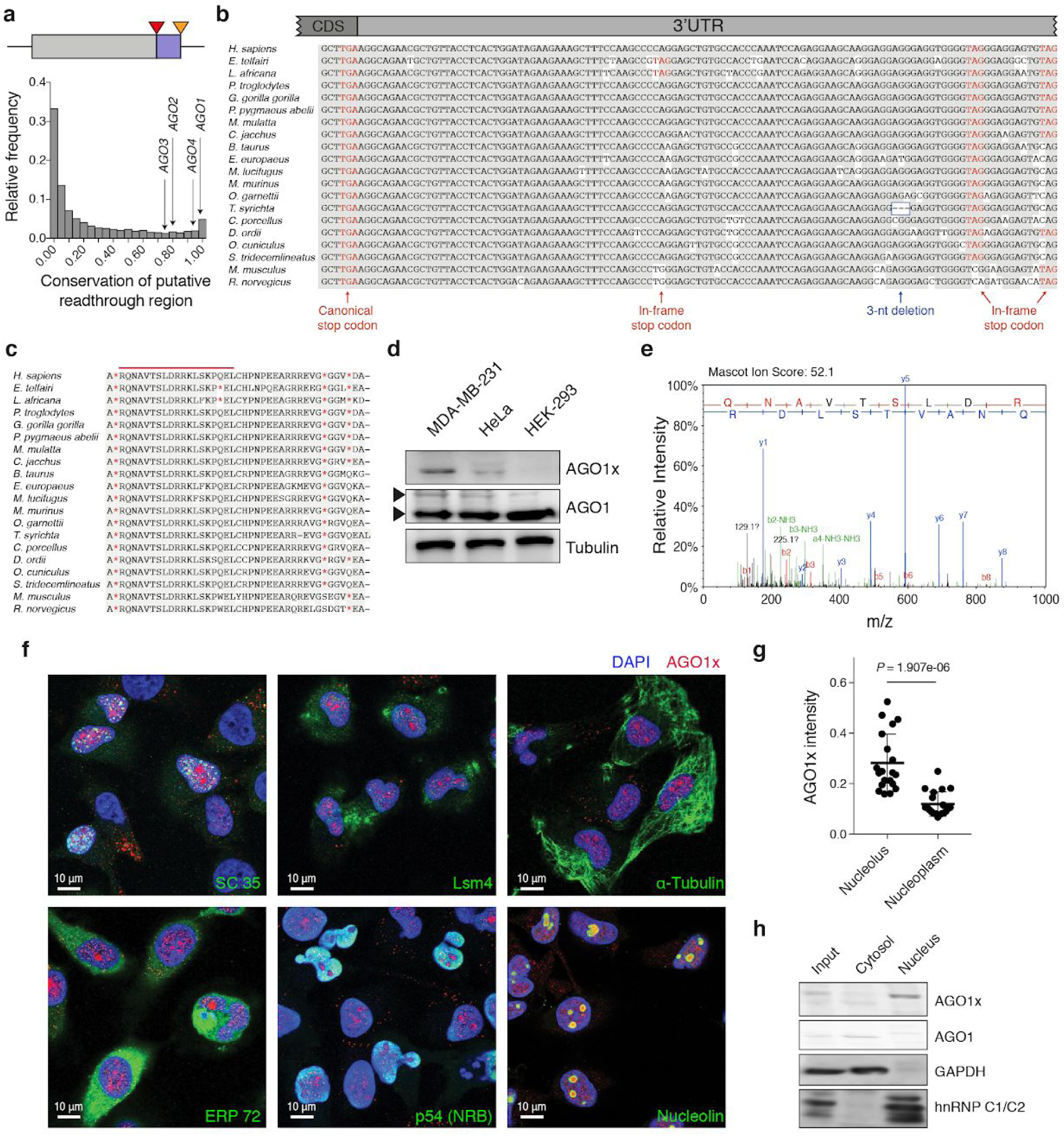
The AGO1x TR isoform localizes to the nucleus. **a**, Histogram of average PhastCons conservation scores (x-axis) for putative TR regions (purple), located downstream of the annotated open reading frame (grey), between the annotated stop codon (red triangle) and the next in-frame stop codon (orange triangle) of all RefSeq-annotated transcripts. The scores of the four members of the human Argonaute protein family are highlighted. **b, c**, Multiple sequence alignment of the region of putative TR in the *AGO1* transcript (**b**) and of the corresponding predicted amino acid sequence (**c**) across vertebrates. The unique peptide targeted by the polyclonal antibody is indicated by the red line. **d**, Western blot showing AGO1x expression in three cell lines. For comparison, a parallel blot was probed with an antibody directed against canonical AGO1. Tubulin staining served as control. **e**, Annotated MS/MS spectrum of the top scoring endogenous peptide “QNAVTSLDR”, specific for AGO1x. The Mascot Ion Score as well as the annotated fragments (blue = y-ions; red = b-ions) together with the corresponding amino acids are indicated. **f**, Representative immunofluorescence images showing the subcellular distribution of AGO1x (red) relative to nuclear and cytoplasmic markers (green). DAPI was used to mark the nucleus (blue). **g**, Mean (+/- s.d.) pixel intensities of AGO1x staining in nucleolus and nucleoplasm, computed from z-stack images of MDA-MB-231 cells (n=20). The P-value was determined using a paired two-tailed t-test. **h**, Representative AGO1x blot from MDA-MB-231 cell fractions. For comparison, AGO1 was also blotted in parallel. GAPDH and hnRNP C1/C2 served as makers of purity of the individual fractions and of relative protein levels.

The expression level of *AGO1* mRNA varies relatively little across normal tissues ^14^ (**Fig. S1a**). We therefore wondered whether larger variation in *AGO1* levels and possibly translational readthrough could arise in cancers as a result of copy number variations of the *AGO1* locus. Indeed, by querying the cbio Cancer Genomics portal ^15^ we found that the *AGO1* locus is amplified with high frequency in ovarian cancer and breast cancer xenografts (**Fig. S1**). An important role for AGO1 in breast cancer has also been reported in a previous study ^16^. Thus, we investigated AGO1x expression in breast cancer and other model cell lines, by western blotting. The commercially available AGO1 antibody consistently identified a cryptic second band of higher molecular weight and lower intensity compared to the band corresponding to the primary signal, in breast cancer cell lines as well as in HeLa cells (**Fig. 1d** and **S2a**). Surmising that this band corresponds to AGO1x, we then obtained a polyclonal antibody directed to the peptide predicted from the readthrough region (**Fig. 1c**, red line). To thoroughly characterize the specificity of this antibody, we overexpressed FLAG-tagged variants of either AGO1 or AGO1x in the MDA-MB-231 breast cancer cell line, and in a complementary experiment, we depleted the *AGO1* transcript with an siRNA pool. Analysis of these cells by western blotting and immunofluorescence demonstrated that our polyclonal AGO1x antibody specifically detected the AGO1x variant, while the commercial AGO1 antibody detected both isoforms (**Fig. S2 b-f**). As we obtained the strongest AGO1x signal in the MDA-MB-231 cell line (**Fig. 1d**), derived from the metastatic site of a human breast tumor, we used primarily this cell line to characterize AGO1x function. HeLa cells, with lower AGO1x expression, were used for additional validation. To further confirm that AGO1x in present in cell lysates, we carried out targeted LC-MS analysis and identified with very high confidence the QNAVTSLDR peptide predicted from the readthrough region (**Fig. 1e**). A translated BLAST ^17^ search against the reference human genome using the NCBI server further showed that this peptide is unique to AGO1x and cannot be derived from any other region of the genome. Furthermore, analysis of RNA sequencing data from our cell lines (see below, **Fig. S5a**) did not identify a single read that could indicate bypass of the annotated stop codon of *AGO1* by alternative splicing. Thus, AGO1x is generated in human breast cancer cell lines, in all likelihood through TR. To gain insight into potential functions of AGO1x, we then investigated its localization in cells, by co-immunofluorescence analysis of AGO1x with several nuclear and cytoplasmic markers. This revealed a striking enrichment of AGO1x in the nuclear compartment of both MDA-MB-231 and HeLa cells, with very little signal from the cytoplasm (**Fig. 1f** and **Fig. S3a**). A more detailed analysis of the nuclear compartment indicated that AGO1x is enriched around nucleoli, as shown by its partial co-localization with the nucleolar marker Nucleolin (**Fig. 1g** and **Fig. S3b, c**). Western blots from nuclear and cytoplasmic fractions of MDA-MB-231 cells corroborated our initial observation that AGO1x is located primarily in the nucleus while the shorter and more abundant canonical isoform is essentially cytoplasmic (**Fig. 1h**).

We next used the CRISPR/CAS9 genome editing tool to investigate the consequences of losing AGO1x expression. To identify robust on-target effects we started from two parental cell lines, MDA-MB-231 and HeLa, and from each, we generated by means of two distinct single guide RNAs (sgRNAs) two derived lines carrying small deletions in the predicted readthrough region (**Fig. 2a**, top panel). By treating the parental cells with an sgRNA designed to target GFP, we also generated a control line. We validated the successful targeting by western blotting, which showed the loss of AGO1x expression, with no apparent effect on the canonical AGO1 isoform in both mutant MDA-MB-231 cell lines (**Fig. 2a**, bottom panel). In culture, the mutant cell lines exhibited a marked reduction in growth relative to the parental line. This was documented both by microscopy at two different time points after seeding equal numbers of cells from the different lines (**Fig. 2b**), as well as through a noninvasive, electrical impedance-based quantification of real-time cell growth ^18^ (**Fig. 2c**). Mutant HeLa lines generated by CRISPR/CAS9 genome editing with the same sgRNAs exhibited similar growth defects (**Fig. S4a-c**). Furthermore, kinetic measurements revealed that beside their slower growth, mutant cell lines also had a reduced migration potential (**Fig. 2d**). This prompted us to evaluate their anchorage-independent growth (**Fig. 2e, S4d**) and self-renewal/differentiation capacities (**Fig. 2f, S4e**) *in vitro*, using soft agar colony and sphere formation assays, respectively. In comparison to control MDA-MB-231 cells, mutant cell lines showed strong impairment in both assays, further underscoring that AGO1x affects cell growth. To determine whether AGO1x may play a role in pathological conditions associated with cell growth, we examined breast cancer patient-derived, paraffin-embedded tissue sections by immunohistochemistry. We found that AGO1x was indeed detectable in the cell nuclei in these samples as well (**Fig. 2g**), its expression being strongly correlated with the proliferative index of the tumours, defined by parallel staining of Ki-67 (**Fig. 2g, h**).

**Figure 2.**
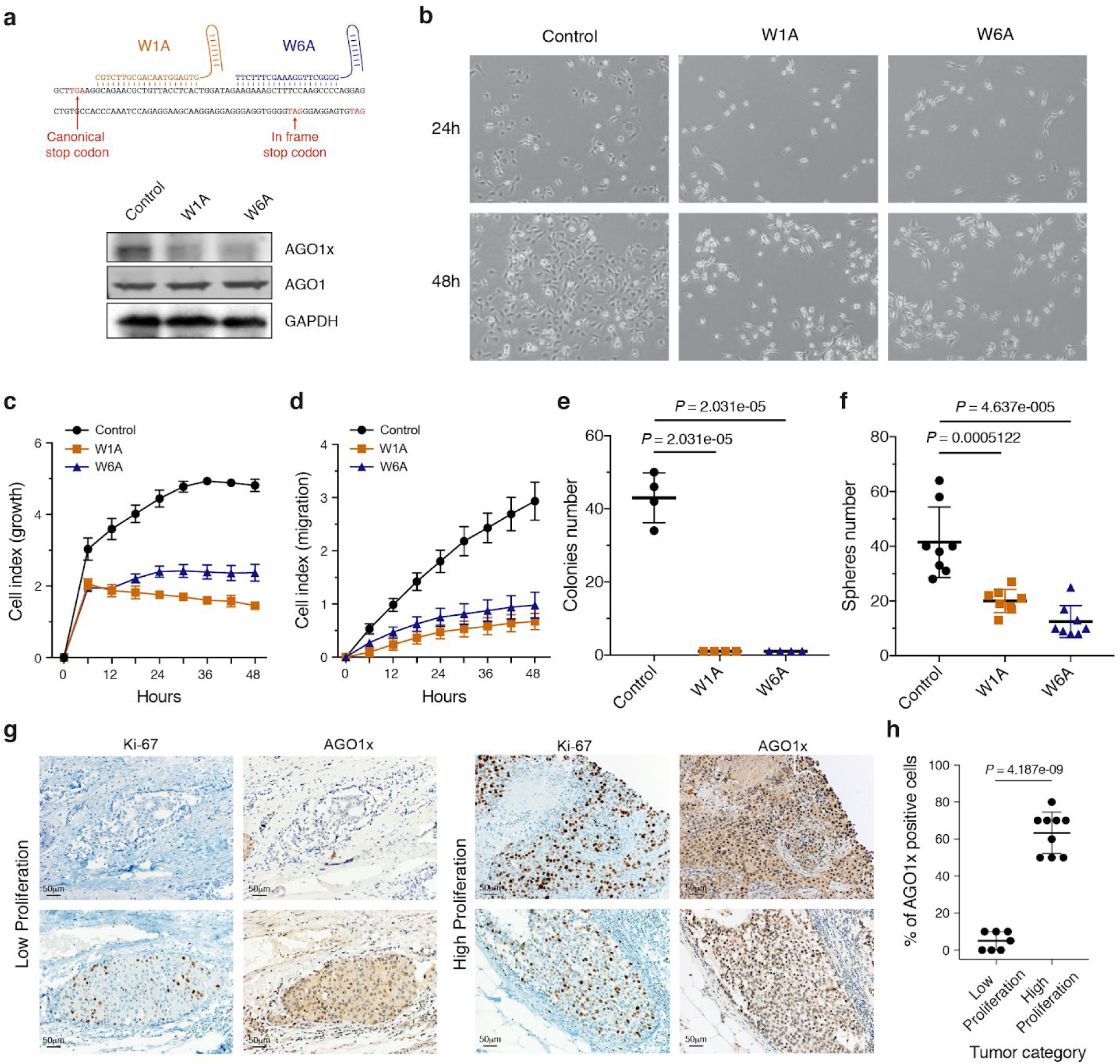
AGO1x promotes cell proliferation. **a**, Scheme of sgRNAs used to generate AGO1x mutant cell lines (top). Western analysis of lysates from mutant MDA-MB-231 cells revealed loss of AGO1x and unchanged levels of canonical AGO1 (bottom). GAPDH was used as control. **b**, Phase contrast images of control and mutant cell lines at 24 and 48 hr after plating equal numbers of cells in individual wells of a six well plate. **c**, Impedance-derived mean (+/- s.d.) cell indices at the indicated time points after seeding equal numbers of control (n=6), W1A (n=5) and W6A (n=5) cells. From 6 hours on, there is a statistical significant difference between control and the two mutant cell lines (*P* < 0.001, two tailed t-test). **d**, Impedance-derived mean (+/- s.d.) cell indices as a function of time for control (n=3), W1A (n=4) and W6A (n=3) cells grown in electronically monitored Boyden chambers. From 12 hours on, there is a statistical significant difference between control and the two mutants (*P* < 0.005, two tailed t-test). **e**, Mean (+/- s.d.) number of colonies obtained from a soft agar colony formation assay of control, W1A and W6A mutant cell lines (n=4). P-value for comparing mutant cell lines to control was determined using an unpaired two-tailed t-test. **f**, Mean (+/- s.d.) number of spheres obtained from a spheres formation assay of control, W1A and W6A mutant cell lines (n=8). P-values computed as in **e**. **g**, AGO1x staining of tissue sections from low-proliferating or high-proliferating breast cancers, classified based on the Ki-67 staining. **h**, Mean (+/- s.d.) percentage of AGO1x positive cells in breast tumors with low (n=7) and high (n=9) proliferation index assessed by the expression level of Ki-67. P-value was determined using an unpaired two-tailed t-test.

To elucidate the mechanism that underlies the impaired cell growth associated with loss of AGO1x expression, we profiled the transcriptomes of the control and the two MDA-MB-231 mutant cell lines. The RNA-seq reads that mapped to the region of readthrough revealed different nucleotide deletions in the mutated cell lines, confirming the successful targeting of this locus (**Fig. S5b**). Several hundred genes had significantly different expression levels (|fold-change| > 2-fold and FDR < 0.01, **Fig. S5c, d**) in mutant compared to control cells. Importantly, the changes were in very strong agreement between the mutant cell lines (*R*=0.92, *P*<0.001, **Fig. 3a**), indicating that they were the result of *AGO1x* targeting rather than to an off-target effect. The expression level of genes related to the interferon alpha response and apoptosis pathways increased in mutant cell lines relative to control, whereas expression of genes involved in cell cycle progression and encoding ribosomal proteins mostly decreased (**Fig. S5e**). Analysis of transcriptional networks underlying the observed changes in gene expression with the ISMARA method ^19^ revealed that the activity of STAT1 and IRF1/2 transcriptional regulators was higher in the mutant cell lines (**Fig. 3b**), consistent with the upregulation of genes from the interferon pathway ^20^ at both transcript (**Fig. 3c**) and protein levels (**Fig. 3d**). Similar to the cellular phenotypes, these findings were reproduced in HeLa cells lines in which the AGO1x readthrough region was genetically targeted (**Fig. S6**). In further agreement with the molecular findings, we documented an increased frequency of apoptosis in the mutant cell lines compared to the control (**Fig. 3e**). Importantly, the growth defect was rescued by ruxolitinib, an inhibitor of the interferon response, implying that the interferon response was the primary reason underlying the impaired growth of mutant lines (**Fig. 3f, g**).

**Figure 3.**
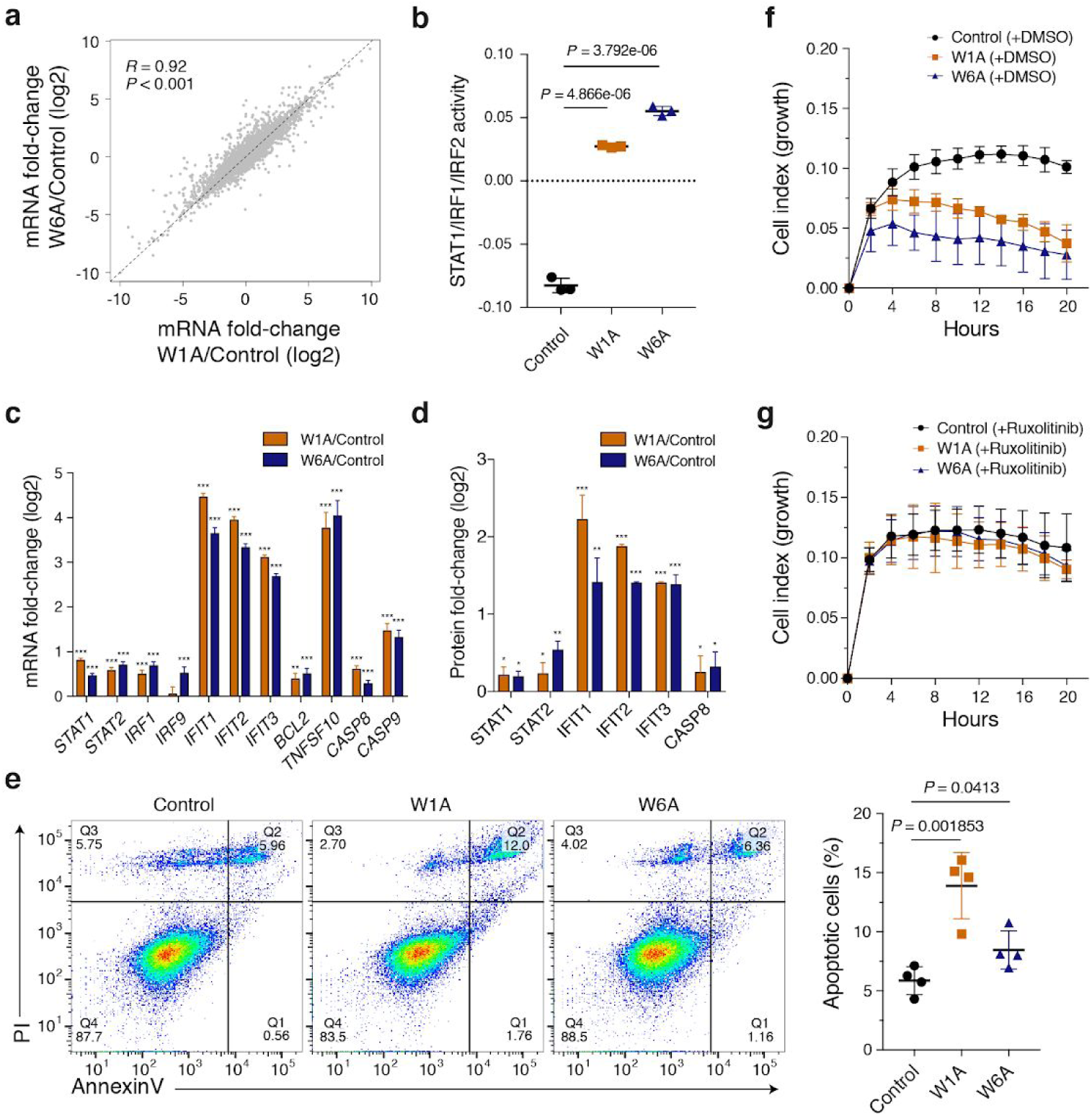
Deletion of AGO1x leads to increased interferon response and apoptosis. **a**, Scatter plot of mRNA fold-changes (log2) in the two mutant cell lines versus control. Shown is also the Pearson correlation coefficient and respective P-value. The dashed line indicates equal fold-changes in the two mutant lines. **b**, Mean (+/- s.d.) activity of STAT1/IRF1/IRF2 transcription factor motifs estimated by ISMARA ^19^ (n=3). Shown is also the P-value in an unpaired two-tailed t-test. **c**, Mean (+/- s.e.m.) mRNA expression fold-changes (log2) of genes involved in the interferon alpha response and apoptosis in the two mutant cell lines relative to control (n=3). **d**, Mean (+/- s.d.) expression fold-changes of the corresponding proteins (if detected in the proteomics data) in the two mutant (W1A, n=2; W6A, n=1) cell lines compared to Control (n=2). Multiple testing corrected P-values for Wald tests (transcriptomics) or unpaired two-tailed t-tests (proteomics) comparing fold-changes with respect to control are depicted above each bar (* *P* < 0.05, ** *P* < 0.01, *** *P* < 0.001). **e**, Representative result of the apoptosis assay using AnnexinV and propidium iodide (PI) staining in the control and the two mutant cell lines (left). The percentage of cells in each quadrant is depicted for each cell line. Quantification of the mean (+/- s.d.) percentage of apoptotic cells (Q1+Q2, AnnexinV+) across the different cell lines (n=4) (right). Shown is the P-value determined by the unpaired two-tailed t-test. **f, g**, Impedance-derived mean (+/- s.d.) cell index values at the indicated time points of growth after seeding equal numbers of control, W1A and W6A cells, upon treatment with DMSO (**f**) or Ruxolitinib (**g**). For the DMSO treatment: Control (n=4), W1A (n=3) and W6A (n=3). For the Ruxolitinib treatment: Control (n=6), W1A (n=3) and W6A (n=4). From 6 hours on, there is a statistical significant difference between control and the two mutants after DMSO treatment (*P* < 0.05, two tailed t-test), whereas no statistically significant difference is found after Ruxolitinib treatment.

Hypothesizing that the activation of the interferon response observed upon AGO1x inhibition is due to the accumulation of double-stranded RNAs (dsRNAs) ^21,22^, we examined the levels of intracellular dsRNA sensors ^20^. We found that their expression was indeed increased, both at mRNA and protein levels (**Fig. 4a, b**). In addition, we uncovered evidence for protein phosphorylation resulting from downstream interferon signaling (**Fig. 4c**). Staining by SCICONS J2 antibody ^23^ confirmed the accumulation of dsRNAs in AGO1x mutant lines relative to control (**Fig. 4d**), while sequencing of RNAs isolated by J2 antibody-based pulldown revealed a strong enrichment of rRNAs (**Fig. 4e**). The rRNAs and mRNAs that accumulated in the mutant lines had a higher proportion of G/C nucleotides and thus a higher propensity of forming stable secondary structures in comparison to RNAs that were not pulled down by the J2 antibody (**Fig. 4f, g**). We confirmed the direct AGO1x interaction with RNAs identified in the J2 antibody-based pulldown by carrying out an additional AGO1x IP followed by quantitative PCR (**Fig. 4h**). Mass spectrometry analysis identified only very few proteins that were reproducibly co-immunoprecipitated from nuclear extracts by AGO1x-specific antibody. Consistent with our microscopic observations of peri-nucleolar localization of AGO1x, the prominent nucleolar component fibrillarin was highly enriched in the AGO1x-IP. Two other AGO1x partners identified in this experiment were the Polyribonucleotide Nucleotidyltransferase 1 (PNPT1) and the DExH-Box helicase 9 (DHX9) (**Fig. 4i**), two proteins previously linked to dsRNAs ^24,25^. The interaction of AGO1x with these proteins was further confirmed by western blotting (**Fig. 4j**). Further indicating its participation in dsRNA processing, the siRNA-mediated depletion of PNPT1 exacerbated the accumulation of dsRNAs in the AGO1x mutant cell line (**Fig. 4k**).

**Figure 4.**
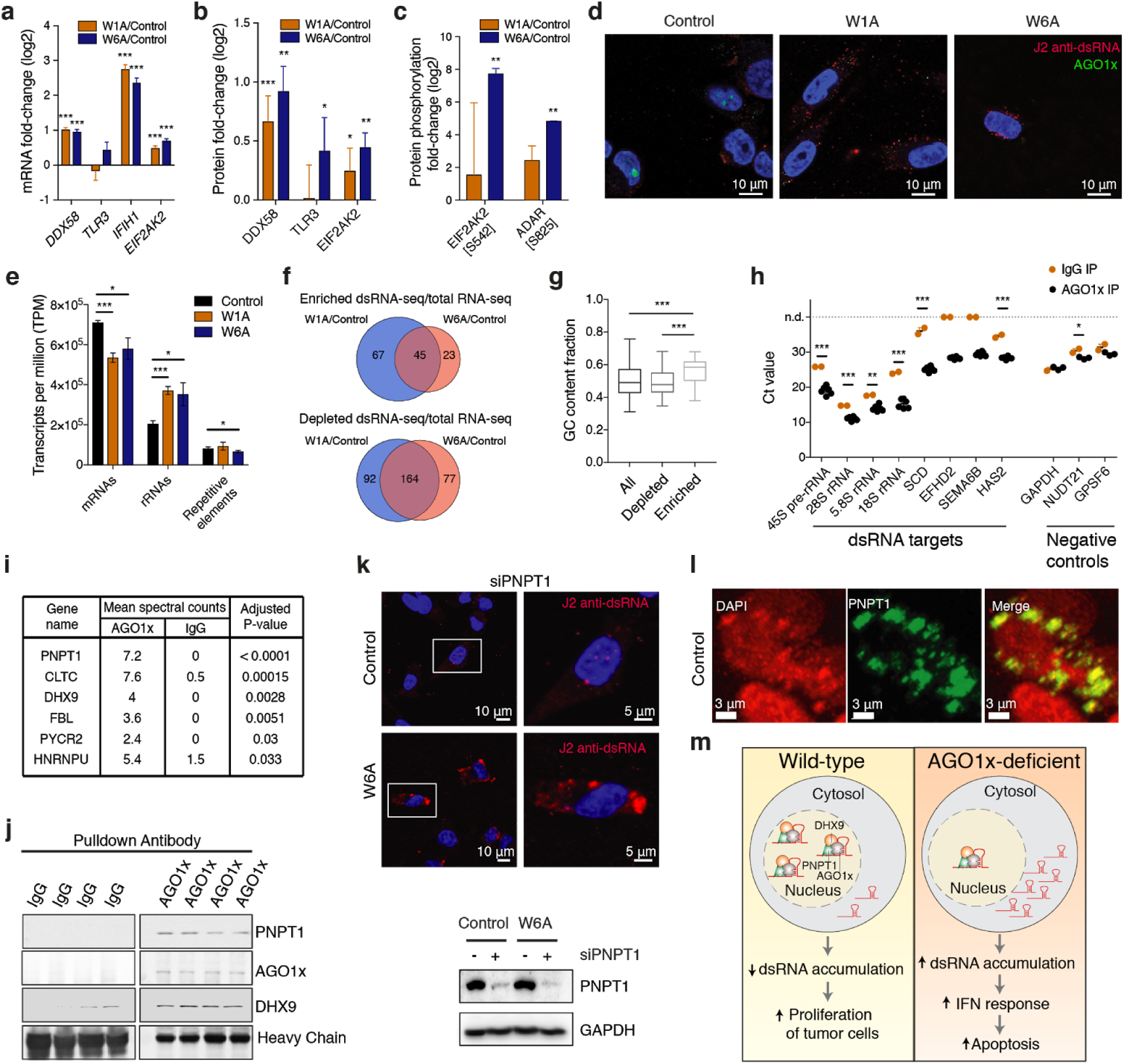
AGO1x interacts with PNPT1 and DHX9 to prevent dsRNA accumulation. **a**, Mean (+/- s.e.m.) mRNA fold-changes (log2) of dsRNAs sensors from the interferon response pathway in the two mutant cell lines compared to control (n=3). **b**, Mean (+/- s.d.) protein fold-changes for the genes from (**a**) that were detected in the isobaric mass tagged based proteomics dataset in the two mutant (W1A, n=2; W6A, n=1) lines compared to control (n=2). **c**, Mean (+/- s.d.) fold-changes in phosphorylated protein levels derived from enrichment of phosphopeptides in the two mutant lines (W1A, n=2; W6A, n=2) compared to control (n=1). Multiple test corrected P-values for Wald tests (transcriptomics) or unpaired two-tailed t-tests (proteomics) comparing fold-changes with respect to control are depicted above each bar (* *P* < 0.05, ** *P* < 0.01, *** *P* < 0.001). **d**, Representative immunofluorescence images of control and mutant MDA-MB-231 cells stained with AGO1x (green) and J2 antibody (red). DAPI was used to mark the nucleus (blue). **e**, Relative abundance of various RNA species in J2 antibody immunoprecipitates from control and mutant cell lines. Multiple testing corrected P-values from unpaired two-tailed t-tests are depicted above each comparison (* *P* < 0.05, ** *P* < 0.01, *** *P* < 0.001). **f**, Venn diagrams showing the intersection between RNA transcripts that were consistently enriched (top) or depleted (bottom) in the dsRNA-seq relative to total RNA-seq in the two mutant cell lines compared to control (n=3). **g**, Boxplots showing the proportion of G/C nucleotides in all genes and in genes depleted/enriched in dsRNA-seq compared to total RNA-seq. Shown are the P-values in the non-parametric Mann-Whitney U test (* *P* < 0.05, ** *P* < 0.01, *** *P* < 0.001). Boxes extend from the 25th to 75th percentiles (inter-quartile range (IQR)), horizontal lines represent the median, whiskers indicate the lowest and highest datum within 1.5*IQR from the lower and upper quartiles, respectively. **h**, Quantification of transcript abundance by qRT-PCR in AGO1x (black, n=6 for putative targets, n=3 for negative controls) or IgG (orange, n=2) IP from control cells. n.d., not detected. Multiple test corrected P-values for a unpaired two-tailed t-test are depicted above each comparison (* *P* < 0.05, ** *P* < 0.01, *** *P* < 0.001). **i**, Summary of mass-spectrometric analysis of AGO1x (n=5) or IgG (n=2) IP from control cells. P-values were calculated using Fisher’s exact test and corrected for multiple testing using the Benjamini-Hochberg method. **j**, Western blot analysis demonstrating the interaction of AGO1x with PNPT1 and DHX9 in control cells. **k**, Representative immunofluorescence images of siPNPT1-treated control and W6A mutant MDA-MB-231 cells stained with J2 antibody (red) (top). DAPI was used to mark the nucleus (blue). The panels on the right show a magnification of the cells in the white boxes depicted on the left panels. Western blot analysis quantifying the depletion of PNPT1 upon siRNA treatment (bottom). **l**, IF images to show overlap of PNPT1 signal with the nucleus. DAPI-stained nuclei are marked in red **m**, Model of AGO1x function. A complex of AGO1x with PNTP1 and DHX9 interacts with nuclear RNAs to prevent the accumulation of double stranded structures, supporting proliferation of breast cancer cells. Depletion of AGO1x leads to deleterious accumulation of dsRNAs, which in turn lead to activation of interferon (IFN) response and apoptosis.

Thus, our study shows that, contrasting with the miRNA-dependent gene silencing function of other Argonaute proteins ^26^, AGO1x interacts with PNPT1 and DHX9 to prevent the inadvertent accumulation of endogenous dsRNAs, which can impair gene expression and trigger the interferon response (**Fig. 4m**). AGO1x interacts directly with rRNAs and some mRNA that are G/C-rich and have the propensity to form stable secondary structures. Consistently, we observed that AGO1x accumulates in peri-nucleolar regions and co-immunoprecipitates with fibrillarin. Two other proteins specifically co-immunoprecipitating with AGO1x, PNPT1 and DHX9, have been previously associated with processes involving dsRNA. Knockdown of the DHX9 helicase was reported to impair rRNA biogenesis ^27^, while very recently, the resolution of dsRNA structures formed by inverted *Alu* repeats by DHX9 was found to promote expression of host transcripts ^24^. PNPT1 is best known for forming an ATP-dependent hSUV3-PNPT1 complex that degrades dsRNA substrates in mitochondria ^25^, although functions of PNPT1 beyond the mitochondria have also been reported ^28,29^. Similar to the latter studies, in our system, PNPT1 co-immunoprecipitated with AGO1x from nuclear extracts (**Fig. 4i**), while imaging revealed its presence in close association with nuclei (**Fig. 4l**).

Especially intriguing in our data is the concept that the increased demand for ribosome biogenesis in rapidly dividing cells engages a previously undescribed mechanism for resolving dsRNA structures in rRNAs and other molecules. This mechanism relies on the production of the AGO1x isoform by stop codon readthrough. The strong evolutionary conservation of *AGO1* 3’ untranslated region, that encodes its C-terminal extension, hints to an important physiological role. Although we here focused on breast cancer cells, we observed AGO1x expression in non-malignant human tissues as well (**Fig. S7**). However, AGO1x expression in normal tissues appears to be restricted to a small proportion of cells, whose nature remains to be further defined. Suppression of the interferon response by AGO1x could be particularly important in pathological conditions where the inhibition of this pathway has been associated with poor clinical outcomes ^30,31^. The broad presence of AGO1x in proliferating cancers cells but not in normal human cells may offer a therapeutic strategy to halt the uncontrolled proliferation of cancer cells.

## Methods

### Cell culture, transfections, treatments and common reagents

MDA-MB-231, HEK-293, and HeLa cells were cultured as described before ^32^. Transfections were performed with Lipofectamine RNAimax (Life Technologies). Western blotting was performed as described earlier ^32^ and the HRP-labelled secondary antibodies were developed with SuperSignal™ West Pico PLUS Chemiluminescent Substrate (ThermoFisher Scientific #34580) or with SuperSignal™ West Femto Maximum Sensitivity Substrate (ThermoFisher Scientific #34095) (AGO1x blot in Fig. 1h). All western blot images were documented with Azure c600 Gel documentation system equipped with a 8.3 MP CCD camera. Ruxolitinib (INCB018424) was obtained from Selleckchem (# S1378) and used at a final concentration of 500 nM in DMSO (Sigma #41639). Details of the plasmids used are included in **Table S1**. The strategy for guide RNA cloning and selection of mutants and control was followed from a previously published protocol ^33^. siRNA and sgRNAs used in the study are listed in **Table S1**. Antibodies used for the study are listed in **Table S2**.

### Genome-wide analysis of C-terminal protein extensions

Putative C-terminal extensions were predicted by taking the mRNA region between the annotated stop codon and the next in-frame stop codon of all RefSeq-annotated (on September 2014) transcripts. PhastCons conservation scores across 45 different vertebrate species for each nucleotide in these regions were downloaded from the UCSC genome browser ^34^ and averaged across the entire regions of putative translational readthrough. Multiple alignments of the 45 vertebrates genomes were also retrieved from the UCSC genome browser, and for Figure 1, the 20 sequences with highest homology to AGO1 extended region were re-aligned using ClustalW ^35^.

### Cell Fractionation

Cells were grown in 60mm dishes for 18-24 hrs and snap frozen immediately in liquid Nitrogen. Subsequently the cells were fractionated as described before ^36^. Fraction lysates obtained were loaded in volume equivalents for each fraction and western blots were developed with Pico PLUS Chemiluminescent Substrate except for AGO1x (with Femto Maximum Sensitivity Substrate).

### Immunofluorescence

For immunofluorescence analysis, cells were fixed with 4% paraformaldehyde for 30 min. Subsequently they were permeabilized and blocked for 30 min with 0.1% Triton X-100 (#T8787, Sigma-Aldrich), 10% goat serum (#16210072, Gibco®, Life Technologies), and 1% BSA (#A9647, Sigma-Aldrich) in PBS (#20012-019, Gibco®, Life Technologies). When AGO1 antibodies were used for staining, the blocking buffer was modified to use donkey serum (D9663-10ML) instead of goat serum. Thereafter, the cells were incubated with primary antibodies (1:100 dilution) in the same buffer at desired dilution overnight at 4°C. Secondary anti-rabbit, anti-goat or anti-mouse antibodies labelled either with Alexa Fluor® 488 dye (green), Alexa Fluor® 568 dye (orange), Alexa Fluor® 594 dye (red) or Alexa Fluor® 647 dye (far red) fluorochromes (Molecular Probes) were used at 1:500 dilutions. The cells were mounted on a glass slide with Vectashield DAPI (Vector Laboratories) and cells were mostly observed and documented with a ZEISS point scanning confocal LSM 700 / LSM 800 Inverted microscopes with a PLAN APO 40X (NA=1.3) and 63X (NA=1.4) oil immersion objectives. Z-stack images were captured wherever mentioned in text.

### Quantification of AGO1x subcellular localization from IF images

Quantitative analysis of confocal z-stacks was performed in Matlab/r2016a (http://www.mathworks.com) with the Image Processing Toolbox ^37^ and ImageJ v1.51n ^38^ (http://imagej.nih.gov/ij/). The processing of every z-stack was executed in three steps. First, we detected the location of every nucleus at each layer of the z-stack. For this we created the nuclei projection mask by applying the maximum function to all blue channel (DAPI) images from the z-stack and subsequently segmenting the resulting image. The mask was further applied to each blue channel image of the z-stack, thus defining the area where nuclei are located. Then, if the nucleus was present in this area, it was segmented. Second, we detected the location of every nucleolus at each layer of the z-stack. We applied the nuclei projection mask to all red channel (Nucleolin) images, to remove the noise outside of nuclei. Then for resulting red channel images we applied the same procedure as for nuclei detection. For assigning nucleoli to corresponding nuclei we considered the overlap projection masks of nuclei and nucleoli. Finally, for green (AGO1x) channel images, we collected the intensity of pixels belonging to nucleoli and nucleoplasm, respectively.

### Transcriptome profiling with total RNA-seq

Total RNA was quality-checked on the Bioanalyzer instrument (Agilent Technologies, Santa Clara, CA, USA) using the RNA 6000 Nano Chip (Agilent, Cat# 5067-1511) and quantified by Spectrophotometry using the NanoDrop ND-1000 Instrument (NanoDrop Technologies, Wilmington, DE, USA). 1µg total RNA was used for library preparation with the TruSeq Stranded mRNA Library Prep Kit High Throughput (Cat# RS-122-2103, Illumina, San Diego, CA, USA). Libraries were quality-checked on the Fragment Analyzer (Advanced Analytical, Ames, IA, USA) using the Standard Sensitivity NGS Fragment Analysis Kit (Cat# DNF-473, Advanced Analytical). The average concentration was 128±12 nmol/L. Samples were pooled to equal molarity. Each pool was quantified by PicoGreen Fluorometric measurement to be adjusted to 1.8pM and used for clustering on the NextSeq 500 instrument (Illumina). Samples were sequenced using the NextSeq 500 High Output Kit 75-cycles (Illumina, Cat# FC-404-1005). Primary data analysis was performed with the Illumina RTA version 2.4.11 and base calling software version bcl2fastq-2.20.0.422.

### Total RNA-seq analysis

Total RNA-seq reads were subjected to 3′ adapter trimming (5’-TGGAATTCTCGGGTGCCAAGG-3’) and quality control (reads shorter than 20 nucleotides or for which over 10% of the nucleotides had a PHRED quality score <20, were discarded). Filtered reads were then mapped to the human transcriptome based on genome assembly hg19 and transcript annotations from RefSeq with the segemehl software ^39^, v0.1.7-411, allowing a minimum mapping accuracy of 90%. Transcript counts were calculated based on uniquely mapped reads and used for differential expression analysis with DESeq2 ^40^.

For the splicing analysis, filtered reads were mapped to the human genome (hg19) with STAR ^41^, v2.6.0c, using default parameters. Read alignments to the transcriptome were visualized using IGV ^42^, v2.4.16.

### dsRNA pulldown and library preparation

dsRNAs were enriched using J2 antibody (Scicons # 10010200) from cell lysates as performed earlier ^43^. The library preparation started from 5ng of RNA based on Fluorometric assessment for each sample. No other selection (poly(A)+ or ribo-depletion) was performed to allow unbiased detection of rRNAs as well as mRNAs. Standard fragmentation / priming step were done as before (for total RNA-seq) to obtain cDNA libraries.

### dsRNA-seq differential expression analysis

RNA-seq reads were subjected to 3′ adapter trimming (5’-GATCGGAAGAGCACACGTCTGAACTCCAGTCAC-3’) and quality control (reads shorter than 20 nucleotides or for which over 10% of the nucleotides had a PHRED quality score <20, were discarded). Filtered reads were then mapped to the human transcriptome based on genome assembly hg19 and transcript annotations from RefSeq with the segemehl software ^39^, v0.1.7-411, allowing a minimum mapping accuracy of 90%. Unmapped reads were additionally mapped to an “artificial transcriptome” composed of consensus sequences for more than 1,000 different repeat elements present in the human genome (including ancestral shared) as defined by the Genetic Information Research Institute Repbase v23.09 (https://www.girinst.org/repbase/). Transcript counts were calculated based on uniquely mapped reads (for mRNAs, rRNAs and repetitive elements) and multi-mapped reads (for repetitive elements), and used for differential expression analysis with DESeq2 ^40^.

### Comparison between dsRNA-seq and total RNA-seq

To identify RNA transcripts enriched or depleted in the dsRNA pulldown sample, we compare the fold-changes in transcript abundances between the mutant and control cell lines in the dsRNA-seq samples to that observed for total RNA-seq samples. Since fold-changes computed from the different datasets were well correlated, we defined RNA transcripts as being enriched or depleted in dsRNA structures as those that deviated significantly from the linear regression of dsRNA-seq as a function of total RNA-seq fold-changes (|standardized residuals| > 2.5 s.d.). Only RNA transcripts consistently enriched/depleted in the comparisons between the two different mutant cell lines and the control were considered for further analysis.

### qRT-PCR to estimate the abundance of messenger and ribosomal RNAs

50ng of AGO1x-IPed RNA was used for reverse transcription following the manufacturer’s protocol and cycling conditions (High-Capacity cDNA Reverse Transcription Kit, Thermo Fisher Scientific). Subsequently, the RT reaction was diluted 4 fold with water and subjected to q-PCR in a 96 well format, using primers specific to individual genes and GoTaq® qPCR Master Mix (Promega). The incubation and cycling conditions were set as described in the kit and the plates were analysed in a StepOnePlus Real-Time PCR System (Thermo Scientific).

### Identification of AGO1x by targeted LC-MS

For each sample, 5×10^6^ cells were lysed and AGO1x-affinity purified using the specific antibody. After washing, the beads and the associated proteins were reduced with 5 mM TCEP for 30 min at 60°C and alkylated with 10 mM chloroacetamide for 30 min at 37 °C. Subsequently, the protein sample was digested by incubation with sequencing-grade modified trypsin (1/50, w/w; Promega, Madison, Wisconsin) overnight at 37°C. Finally, peptides were desalted on C18 reversed phase spin columns according to the manufacturer’s instructions (Microspin, Harvard Apparatus), dried under vacuum and stored at −80°C until further processing. Next, 0.1 µg of peptides of each sample were subjected to targeted MS analysis. Therefore, 6 peptide sequences specific for AGO1 and AGO1x were selected and imported into the Skyline software V2.1, https://brendanx-uw1.gs.washington.edu/labkey/project/home/software/Skyline/begin.view).

Then, a mass isolation list comprising the precursor ion masses with charge 2 and 3+ of all peptides was exported and used for parallel reaction monitoring (PRM) ^44^, quantification on a Q-Exactive HF platform. In brief, peptide separation was carried out using an EASY nLC-1000 system (Thermo Fisher Scientific) equipped with a RP-HPLC column (75μm × 30cm) packed in-house with C18 resin (ReproSil-Pur C18–AQ, 1.9μm resin; Dr. Maisch GmbH, Ammerbuch-Entringen, Germany) using a linear gradient from 95% solvent A (0.1% formic acid, 2% acetonitrile) and 5% solvent B (98% acetonitrile, 0.1% formic acid) to 45% solvent B over 60 min at a flow rate of 0.2μl/min. 3×10^6^ ions were accumulated for MS1 and MS2 and scanned at a resolution of 60,000 FWHM (at 200 m/z). Fill time was set to 150 ms for both scan types. For MS2, a normalized collision energy of 28% was employed, the ion isolation window was set to 0.4 Th and the first mass was fixed to 100 Th. All raw-files were imported into Skyline for protein/peptide quantification. To control for variation in injected sample amounts, samples were normalized using the total ion current from the MS1 scans. Finally, all generated raw files were subjected to standard database searching to validate the peptide identity. Therefore, the acquired raw-files were converted to the Mascot generic file (mgf) format using the msconvert tool (part of ProteoWizard, version 3.0.4624 (2013-6-3)). Using the Mascot algorithm (Matrix Science, Version 2.4.0), the mgf files were searched against a decoy database containing normal and reverse sequences of the predicted SwissProt entries of Homo sapiens (www.uniprot.org, release date 29/06/2015), the C-terminal extension in AGO1x and commonly observed contaminants (in total 41,159 protein sequences) generated using the SequenceReverser tool from the MaxQuant software (Version 1.0.13.13). The precursor ion tolerance was set to 10 ppm and fragment ion tolerance was set to 0.02 Da. The search criteria were set as follows: full tryptic specificity was required (cleavage after arginine residues unless followed by proline), 1 missed cleavage was allowed, carbamidomethylation (C), was set as fixed modification and oxidation (M) was set as variable modifications. Next, the database search results were imported to the Scaffold Q+ software (version 4.3.3, Proteome Software Inc., Portland, OR) and the peptide and protein false identification rate was set to 1% based on the number of decoy hits.

### Global proteome and phosphoproteome analysis by shotgun LC-MS

For each sample, 5×10^6^ cells were washed twice with ice-cold 1x phosphate-buffered saline (PBS) and lysed in 100 μl urea lysis buffer (8 M urea (AppliChem), 0.1 M Ammonium Bicarbonate (Sigma), 1× PhosSTOP (Roche)). Samples were vortexed, sonicated at 4°C (Hielscher), shaked for 5 min on a thermomixer (Eppendorf) at room temperature and centrifuged for 20 min at 4°C full speed. Supernatants were collected and protein concentration was measured with BCA Protein Assay kit (Invitrogen). Per sample, a total of 300 µg of protein mass were reduced with tris(2-carboxyethyl)phosphine (TCEP) at a final concentration of 10 mM at 37°C for 1 hour, alkylated with 20 mM chloroacetamide (CAM, Sigma) at 37°C for 30 minutes and incubated for 4 h with Lys-C endopeptidase (1:200 w/w). After diluting samples with 0.1 M Ammonium Bicarbonate to a final urea concentration of 1.6 M, proteins were further digested overnight at 37°C with sequencing-grade modified trypsin (Promega) at a protein-to-enzyme ratio of 50:1. Subsequently, peptides were desalted on a C18 Sep-Pak cartridge (VAC 3cc, 500 mg, Waters) according to the manufacturer’s instructions, split in peptide aliquots of 200 and 25 µg, dried under vacuum and stored at −80°C until further use.

For proteome profiling, sample aliquots containing 25 µg of dried peptides were subsequently labeled with isobaric tag (TMT 6-plex, Thermo Fisher Scientific) following a recently established protocol ^45^. To control for ratio distortion during quantification, a peptide calibration mixture consisting of six digested standard proteins mixed in different amounts were added to each sample before TMT labeling. After pooling the TMT labeled peptide samples, peptides were again desalted on C18 reversed-phase spin columns according to the manufacturer’s instructions (Macrospin, Harvard Apparatus) and dried under vacuum. TMT-labeled peptides were fractionated by high-pH reversed phase separation using a XBridge Peptide BEH C18 column (3,5 µm, 130 Å, 1 mm × 150 mm, Waters) on an Agilent 1260 Infinity HPLC system. Peptides were loaded on column in buffer A (ammonium formate (20 mM, pH 10) in water) and eluted using a two-step linear gradient starting from 2% to 10% in 5 minutes and then to 50% (v/v) buffer B (90% acetonitrile / 10% ammonium formate (20 mM, pH 10) over 55 minutes at a flow rate of 42 µl/min. Elution of peptides was monitored with a UV detector (215 nm, 254 nm). A total of 36 fractions were collected, pooled into 12 fractions using a post-concatenation strategy as previously described ^46^, dried under vacuum and subjected to LC-MS/MS analysis.

For phosphoproteome profiling, sample aliquots containing 200 µg of dried peptides were subjected to phosphopeptide enrichment using IMAC cartridges and a BRAVO AssayMAP liquid handling platform (Agilent) as recently described ^47^.

The setup of the μRPLC-MS system was as described previously ^45^. Chromatographic separation of peptides was carried out using an EASY nano-LC 1000 system (Thermo Fisher Scientific), equipped with a heated RP-HPLC column (75 μm × 30 cm) packed in-house with 1.9 μm C18 resin (Reprosil-AQ Pur, Dr. Maisch). Aliquots of 1 μg total peptides were analyzed per LC-MS/MS run using a linear gradient ranging from 95% solvent A (0.15% formic acid, 2% acetonitrile) and 5% solvent B (98% acetonitrile, 2% water, 0.15% formic acid) to 30% solvent B over 90 minutes at a flow rate of 200 nl/min. Mass spectrometry analysis was performed on a Q-Exactive HF mass spectrometer equipped with a nanoelectrospray ion source (both Thermo Fisher Scientific) and a custom made column heater set to 60°C. 3E6 ions were collected for MS1 scans for no more than 100 ms and analyzed at a resolution of 120,000 FWHM (at 200 m/z). MS2 scans were acquired of the 10 most intense precursor ions at a target setting of 100,000 ions, accumulation time of 50 ms, isolation window of 1.1 Th and at resolution of 30,000 FWHM (at 200 m/z) using a normalized collision energy of 35%. For phosphopeptide enriched samples, the isolation window was set to 1.4 Th and a normalized collision energy of 28% was applied. Total cycle time was approximately 1-2 seconds.

For proteome profiling, the raw data files were processed as described above using the Mascot and Scaffold software and TMT reporter ion intensities were extracted. Phosphopeptide enriched samples were analyzed by label-free quantification. Therefore, the acquired raw-files were imported into the Progenesis QI software (v2.0, Nonlinear Dynamics Limited), which was used to extract peptide precursor ion intensities across all samples applying the default parameters.

Quantitative analysis results from label-free and TMT quantification were further processed using the SafeQuant R package v.2.3.2. (https://github.com/eahrne/SafeQuant/) to obtain protein relative abundances. This analysis included global data normalization by equalizing the total peak/reporter areas across all LC-MS runs, summation of peak areas per protein and LC-MS/MS run, followed by calculation of protein abundance ratios. Only isoform specific peptide ion signals were considered for quantification. The summarized protein expression values were used for statistical testing of between condition differentially abundant proteins. Here, empirical Bayes moderated t-tests were applied, as implemented in the R/Bioconductor limma package (http://bioconductor.org/packages/release/bioc/html/limma.html). The resulting per protein and condition comparison P-values were adjusted for multiple testing using the Benjamini-Hochberg method.

### Identification of AGO1x interacting proteins by IP

For identification of the interactors of AGO1x in the cell, MDA-MB-231 cells were lysed in a two step reaction. In a pre-clearing step, the cells were washed to deplete free cytosolic proteins which would enrich for AGO1x in the pulldown while depleting background noise. This was achieved by incubating the cells in a buffer containing 25mM Tris/HCl, 150 mM KCl, 2mM EDTA, 0.05% NP40, 1 mM NaF, 1 mM DTT supplemented with protease inhibitor cocktail for 5 mins and removing the supernatant after centrifugation. Subsequently the pellets were lysed in a buffer containing 50 mM Tris/HCl, pH 7.5, 150 mM NaCl, 5 mM EDTA, 0.5% NP-40, 10% glycerol, 1 mM NaF, 0.5 mM DTT supplemented with Protease Inhibitor Cocktail-EDTA Free (Roche). The lysates were clarified by spinning at 2000 g and then incubated overnight with 10ug of AGO1x or control IgG antibody. Subsequently, 100 µl of Dynabeads® Protein G (Thermo Scientific) were added to each sample for 4 hrs to facilitate the binding of beads to the antibody. Finally, beads were washed thrice with 50 mM Tris/HCl, pH 7.5, 300 mM NaCl, 5 mM MgCl2, 0.05% (v/v) NP-40 and 1 mM NaF and once with PBS. Proteins were eluted with 50 µl 100 mM glycine pH 2.6 with incubation at room temperature for 5 mins. The eluate was neutralized with 1M NaOH (1:20) and followed up with either Western blotting or LC-MS analysis.

### Identification of AGO1x protein-interactions by shotgun LC-MS

The setup of the μRPLC-MS system was as described above using an EASY nano-LC 1000 system coupled to a LTQ-Orbitrap Elite (both Thermo Fisher Scientific). Peptide separation was performed as described above. Mass spectrometry analysis was performed on a dual pressure LTQ-Orbitrap Elite mass spectrometer equipped with a nanoelectrospray ion source and a custom made column heater set to 60°C. Each MS1 scan (acquired in the Orbitrap) was followed by collision-induced-dissociation (CID, acquired in the linear ion trap) of the 20 most abundant precursor ions with dynamic exclusion for 60 seconds. Total cycle time was approximately 2 s. For MS1, 1E6 ions were accumulated in the Orbitrap cell over a maximum time of 300 ms and scanned at a resolution of 240,000 FWHM (at 400 m/z). MS2 scans were acquired at a target setting of 10,000 ions, accumulation time of 25 ms and rapid scan rate using a normalized collision energy of 35%. The preview mode was activated and the mass selection window was set to 2 Da.

For data analysis, the acquired raw-files were converted to the Mascot generic file (mgf) format using the msconvert tool (part of ProteoWizard, version 3.0.4624 (2013-6-3)). Using the Mascot algorithm (Matrix Science, Version 2.4.0), the mgf files were searched against a decoy database containing normal and reverse sequences of the predicted SwissProt entries of Homo sapiens (www.uniprot.org, release date 29/06/2015), the C-terminal extension in AGO1x and commonly observed contaminants (in total 41,159 protein sequences) generated using the SequenceReverser tool from the MaxQuant software (Version 1.0.13.13). The precursor ion tolerance was set to 10 ppm and fragment ion tolerance was set to 0.02 Da. The search criteria were set as follows: full tryptic specificity was required (cleavage after arginine residues unless followed by proline), 1 missed cleavage was allowed, carbamidomethylation (C), was set as fixed modification and oxidation (M) was set as variable modifications. Next, the database search results were imported to the Scaffold Q+ software (version 4.3.3, Proteome Software Inc., Portland, OR) and the peptide and protein false identification rate was set to 1% based on the number of decoy hits. Peptide spectrum counts were used for differential protein analysis.

### Patient samples

Sixteen consecutive breast cancers tissues and tumor free tissues from six organs (breast, lung, kidney, prostate, stomach and colon) were retrieved from the archive of the Institute of Pathology at the University Hospital Basel (Basel). Samples were anonymized prior to analysis and the approval for the use of these samples has been granted by the local ethics committee (Number: 2016-01748).

### Immunohistochemistry

Immunohistochemical (IHC) staining for AGO1x and Ki-67 was performed on 4 µm sections of FFPE tissue using primary antibodies against anti-AGO1x (Luzerna-Chem; clone RBP 1510, dilution 1:100, citrate buffer pH 6.0 antigen retrieval) and anti-Ki-67 (Dako; clone IR626, dilution 1:200, citrate buffer pH 6.0 antigen retrieval). Staining procedures were performed on Leica Bond III autostainer using Bond ancillary reagents and a Refine Polymer Detection system according to the manufacturer guidelines. Immunoreactivity for AGO1x and Ki-67 was performed semi-quantitatively as the number of positive tumor cells over the total number of tumor as previously described ^48^. All slides were evaluated by a trained pathologist (LMT). Tumors were classified into lowly- or highly-proliferative based on Ki-67 positive cells in accordance to the St. Gallen’s guidelines as previously described ^49^.

### Soft agar colony formation assay

Soft agar assays were carried out in 6-well plates previously coated with a 5 ml layer of medium containing 40% Dulbecco’s modified Eagle’s medium 2X (Thermo Fisher Scientific), 10% FBS (Gibco, Thermo Fisher Scientific), 10% TPB Buffer (Thermo Fisher Scientific) and 0.5% of Noble Agar (Difco). 5×10^4^ cells were diluted in 1.5 ml of 0.16% agar containing medium, layered onto the bed of 0.5% agar in duplicate and left growing for 2 weeks in the incubator. Multiple fields were imaged using an inverted microscope and cell colonies were counted using the ImageJ software.

### Proliferation assay

Cell growth was assayed using the xCELLigence system (RTCA, ACEA Biosciences, San Diego, CA). Background impedance of the xCELLigence system was measured for 12 sec using 50 μL of room temperature cell culture media in each well of E-plate 16. After reaching 75% confluence, the cells were washed with PBS and detached from the flasks using a short treatment with trypsin/EDTA. 10,000 cells were dispensed into each well of an E-plate 16. Growth and proliferation of the cells were monitored every 15 min up to 48 hrs via the incorporated sensor electrode arrays of the xCELLigence system, using the RTCA-integrated software according to the manufacturer’s parameters.

### Migration assay

Migration assay was performed with CIM plates using the xCELLigence system (RTCA, ACEA Biosciences). 3×10^4^ cells were plated in each well according to manufacturer’s instruction and migration was monitored up to 48 hrs after seeding using the RTCA-integrated software, according to the manufacturer’s protocols ^50^.

### Sphere formation assay

10^3^ cells were resuspended in 4 ml of STEM medium containing DMEM/F12 (Thermo Fisher Scientific), B27 supplement (Thermo Fisher Scientific, 1X), EGF (Sigma-Aldrich, 20 ng/ml) and FGF (Sigma-Aldrich, 10 ng/ml). The cells were then plated in in T25 flasks pre-coated with 1% Noble Agar (Difco). Fresh aliquots of medium were added every 3 days and after 10 days the spheres were observed and counted on a Olympus CKX41 inverted microscope equipped with a SC 30 digital camera (Olympus) and counted using the ImageJ software.

### Apoptosis Assay

Steady state apoptosis assays were performed with the Dead Cell Apoptosis Kit with Annexin V Alexa Fluor™ 488 & Propidium Iodide (PI) obtained from ThermoFisher Scientific (#V13241) and assays were performed according to manufacturer’s protocol. Briefly, approx. 10^6^ cells were seeded in a 60 mm cell culture plate and incubated at 37°C in an incubator as normal. Subsequently the cells were harvested with Trypsin EDTA after overnight growth (between 16-18 hrs) and stained with Annexin V conjugated with Alexa 488. For detection of dead/necrotic cells, PI counter staining was also performed. Negative staining controls and single dye staining controls were also made for each cell type for offline gating analyses with FlowJo®. All samples were processed in a BD FACSCanto II analyzer.

### Motif activity analysis

The webserver ISMARA (https://ismara.unibas.ch/) was used to estimate the activities of different transcription factor binding motifs in the different mRNA-seq samples.

### Gene set enrichment analysis

The tool GSEA v2.2.3 (http://software.broadinstitute.org/gsea/index.jsp) was used to calculate the enrichment of gene sets derived from the KEGG pathway database and the Hallmark collection. To estimate significance of the enrichments, the number of permutations was set to 1000 and the permutation type was set to gene sets.

### Statistical analysis

Statistical analysis was performed with Prism 7.0c (GraphPad). P-values were calculated with unpaired two-tailed Student’s t test unless otherwise noted. *P* < 0.05 was considered significant.

### Code availability

Any custom code developed for this study will be made available upon request.

### Data Availability

Sequencing data from this study have been submitted to the Sequence Read Archive under the accession number SRP136692. The MS proteomics data have been deposited to ProteomeXchange with the identifier PXD009401.

## Acknowledgements

We thank Alexandra Gnann for support with cloning, Dr. Noam Stern-Ginossar for suggestions of the interferon pathway experiments, and members of the Zavolan group for comments and suggestions. We would also like to thank Philippe Demougin from the Genomics Facility Basel for genomic library preparation, and the sciCORE team for their maintenance of the HPC facility at the University Basel. This work was funded by the Swiss National Science Foundation NCCR project “RNA & Disease” (grant 51NF40_141735) and by a SystemsX.ch Transitional Postdoctoral Fellowship (grant 51FSP0_157344) and a Novartis University of Basel Excellence Scholarship for Life Sciences to J.C.G.. Additional funding sources include Swiss National Science Foundation (Ambizione grant number PZ00P3_168165) to S.P. and Swiss Cancer League (KFS-3995-08-2016 and KLS-3639-02-2015) to S.P. and L.M.T.. G.M. acknowledges funding by the Deutsche Forschungsgemeinschaft (DFG, SFB960).

## Author Contributions

J.C.G. and M.Z. conceived and supervised the project. S.G. designed and performed most experiments. J.C.G. performed all computational analysis. S.G. and J.C.G. analyzed data and interpreted results. M.L. performed immunohistochemistry and oncogenic related assays. A.S. performed proteomics analysis. A.P.S. generated CRISPR/CAS9 cell lines. A.B. performed image analysis. B.D. and Sh.G. performed western blots and characterization of mutants. A.L.C. provided resources and generated overexpression cell lines. J.D. set up the immunoprecipitation protocol and provided resources. G.M. provided resources and input on the analysis of AGO1x-specific antibody. S.P and L.T. interpreted oncogenic assays and immunohistochemistry stains and provided resources. S.G., J.C.G., and M.Z. wrote the manuscript.

## References

1. Jungreis, I. et al. Evidence of abundant stop codon readthrough in Drosophila and other metazoa. Genome Res. 21, 2096–2113 (2011).

2. Dunn, J. G., Foo, C. K., Belletier, N. G., Gavis, E. R. & Weissman, J. S. Ribosome profiling reveals pervasive and regulated stop codon readthrough in Drosophila melanogaster. Elife 2, e01179 (2013).

3. Eswarappa, S. M. et al. Programmed translational readthrough generates antiangiogenic VEGF-Ax. Cell 157, 1605–1618 (2014).

4. Schueren, F. et al. Peroxisomal lactate dehydrogenase is generated by translational readthrough in mammals. Elife 3, e03640 (2014).

5. Yordanova, M. M. et al. AMD1 mRNA employs ribosome stalling as a mechanism for molecular memory formation. Nature 553, 356–360 (2018).

6. Arribere, J. A. et al. Translation readthrough mitigation. Nature 534, 719–723 (2016).

7. Meister, G. Argonaute proteins: functional insights and emerging roles. Nat. Rev. Genet. 14, 447–459 (2013).

8. Landthaler, M. et al. Molecular characterization of human Argonaute-containing ribonucleoprotein complexes and their bound target mRNAs. RNA 14, 2580–2596 (2008).

9. Ameyar-Zazoua, M. et al. Argonaute proteins couple chromatin silencing to alternative splicing. Nat. Struct. Mol. Biol. 19, 998–1004 (2012).

10. Huang, V. et al. Ago1 Interacts with RNA polymerase II and binds to the promoters of actively transcribed genes in human cancer cells. PLoS Genet. 9, e1003821 (2013).

11. Skourti-Stathaki, K., Kamieniarz-Gdula, K. & Proudfoot, N. J. R-loops induce repressive chromatin marks over mammalian gene terminators. Nature 516, 436–439 (2014).

12. Alló, M. et al. Argonaute-1 binds transcriptional enhancers and controls constitutive and alternative splicing in human cells. Proc. Natl. Acad. Sci. U. S. A. 111, 15622–15629 (2014).

13. Xin, H., Zhong, C., Nudleman, E. & Ferrara, N. Evidence for Pro-angiogenic Functions of VEGF-Ax. Cell 167, 275–284.e6 (2016).

14. Uhlén, M. et al. Proteomics. Tissue-based map of the human proteome. Science 347, 1260419 (2015).

15. Cerami, E. et al. The cBio cancer genomics portal: an open platform for exploring multidimensional cancer genomics data. Cancer Discov. 2, 401–404 (2012).

16. Sung, H. et al. Common genetic polymorphisms of microRNA biogenesis pathway genes and risk of breast cancer: a case-control study in Korea. Breast Cancer Res. Treat. 130, 939–951 (2011).

17. Madden, T. The BLAST Sequence Analysis Tool. (National Center for Biotechnology Information (US), 2013).

18. Atienza, J. M., Zhu, J., Wang, X., Xu, X. & Abassi, Y. Dynamic monitoring of cell adhesion and spreading on microelectronic sensor arrays. J. Biomol. Screen. 10, 795–805 (2005).

19. Balwierz, P. J. et al. ISMARA: automated modeling of genomic signals as a democracy of regulatory motifs. Genome Res. 24, 869–884 (2014).

20. Schneider, W. M., Chevillotte, M. D. & Rice, C. M. Interferon-stimulated genes: a complex web of host defenses. Annu. Rev. Immunol. 32, 513–545 (2014).

21. Field, A. K., Tytell, A. A., Lampson, G. P. & Hilleman, M. R. Inducers of interferon and host resistance. II. Multistranded synthetic polynucleotide complexes. Proc. Natl. Acad. Sci. U. S. A. 58, 1004–1010 (1967).

22. Lampson, G. P., Tytell, A. A., Field, A. K., Nemes, M. M. & Hilleman, M. R. Inducers of interferon and host resistance. I. Double-stranded RNA from extracts of Penicillium funiculosum. Proc. Natl. Acad. Sci. U. S. A. 58, 782–789 (1967).

23. Schönborn, J. et al. Monoclonal antibodies to double-stranded RNA as probes of RNA structure in crude nucleic acid extracts. Nucleic Acids Res. 19, 2993–3000 (1991).

24. Aktaş, T. et al. DHX9 suppresses RNA processing defects originating from the Alu invasion of the human genome. Nature 544, 115–119 (2017).

25. Wang, D. D.-H., Shu, Z., Lieser, S. A., Chen, P.-L. & Lee, W.-H. Human mitochondrial SUV3 and polynucleotide phosphorylase form a 330-kDa heteropentamer to cooperatively degrade double-stranded RNA with a 3’-to-5’ directionality. J. Biol. Chem. 284, 20812–20821 (2009).

26. Leung, A. K. L. The Whereabouts of microRNA Actions: Cytoplasm and Beyond. Trends Cell Biol. 25, 601–610 (2015).

27. Zhang, Y., Forys, J. T., Miceli, A. P., Gwinn, A. S. & Weber, J. D. Identification of DHX33 as a mediator of rRNA synthesis and cell growth. Mol. Cell. Biol. 31, 4676–4691 (2011).

28. Sokhi, U. K. et al. Analysis of Global Changes in Gene Expression Induced by Human Polynucleotide Phosphorylase (hPNPase old-35). J. Cell. Physiol. 229, 1952–1962 (2014).

29. French, S. W. et al. The TCL1 oncoprotein binds the RNase PH domains of the PNPase exoribonuclease without affecting its RNA degrading activity. Cancer Lett. 248, 198–210 (2007).

30. Chiappinelli, K. B. et al. Inhibiting DNA Methylation Causes an Interferon Response in Cancer via dsRNA Including Endogenous Retroviruses. Cell 162, 974–986 (2015).

31. Gao, J. et al. Loss of IFN-γ Pathway Genes in Tumor Cells as a Mechanism of Resistance to Anti-CTLA-4 Therapy. Cell 167, 397–404.e9 (2016).

32. Ghosh, S., Bose, M., Ray, A. & Bhattacharyya, S. N. Polysome arrest restricts miRNA turnover by preventing exosomal export of miRNA in growth-retarded mammalian cells. Mol. Biol. Cell 26, 1072–1083 (2015).

33. Ran, F. A. et al. Genome engineering using the CRISPR-Cas9 system. Nat. Protoc. 8, 2281–2308 (2013).

34. Kent, W. J. et al. The human genome browser at UCSC. Genome Res. 12, 996–1006 (2002).

35. Larkin, M. A. et al. Clustal W and Clustal X version 2.0. Bioinformatics 23, 2947–2948 (2007).

36. Suzuki, K., Bose, P., Leong-Quong, R. Y., Fujita, D. J. & Riabowol, K. REAP: A two minute cell fractionation method. BMC Res. Notes 3, 294 (2010).

37. Gonzalez, R. C., Woods, R. E. & Eddins, S. Digital Image Processing Using MATLAB: Pearson Prentice Hall. Upper Saddle River, New Jersey (2004).

38. Schneider, C. A., Rasband, W. S. & Eliceiri, K. W. NIH Image to ImageJ: 25 years of image analysis. Nat. Methods 9, 671–675 (2012).

39. Hoffmann, S. et al. Fast mapping of short sequences with mismatches, insertions and deletions using index structures. PLoS Comput. Biol. 5, e1000502 (2009).

40. Love, M. I., Huber, W. & Anders, S. Moderated estimation of fold change and dispersion for RNA-seq data with DESeq2. Genome Biol. 15, 550 (2014).

41. Dobin, A. et al. STAR: ultrafast universal RNA-seq aligner. Bioinformatics 29, 15–21 (2013).

42. Robinson, J. T. et al. Integrative genomics viewer. Nat. Biotechnol. 29, 24–26 (2011).

43. Whipple, J. M. et al. Genome-wide profiling of the C. elegans dsRNAome. RNA 21, 786–800 (2015).

44. Peterson, A. C., Russell, J. D., Bailey, D. J., Westphall, M. S. & Coon, J. J. Parallel reaction monitoring for high resolution and high mass accuracy quantitative, targeted proteomics. Mol. Cell. Proteomics 11, 1475–1488 (2012).

45. Ahrné, E. et al. Evaluation and Improvement of Quantification Accuracy in Isobaric Mass Tag-Based Protein Quantification Experiments. J. Proteome Res. (2016). doi:10.1021/acs.jproteome.6b00066

46. Wang, Y. et al. Reversed-phase chromatography with multiple fraction concatenation strategy for proteome profiling of human MCF10A cells. Proteomics 11, 2019–2026 (2011).

47. Post, H. et al. Robust, Sensitive, and Automated Phosphopeptide Enrichment Optimized for Low Sample Amounts Applied to Primary Hippocampal Neurons. J. Proteome Res. 16, 728–737 (2017).

48. Sepe, R. et al. HMGA1 overexpression is associated with a particular subset of human breast carcinomas. J. Clin. Pathol. 69, 117–121 (2016).

49. Piscuoglio, S. et al. The Genomic Landscape of Male Breast Cancers. Clin. Cancer Res. 22, 4045–4056 (2016).

50. Limame, R. et al. Comparative Analysis of Dynamic Cell Viability, Migration and Invasion Assessments by Novel Real-Time Technology and Classic Endpoint Assays. PLoS One 7, e46536 (2012).

51. Eirew, P. et al. Dynamics of genomic clones in breast cancer patient xenografts at single-cell resolution. Nature 518, 422–426 (2015).

52. Cancer Genome Atlas Research Network. Integrated genomic analyses of ovarian carcinoma. Nature 474, 609–615 (2011).

53. Beltran, H. et al. Divergent clonal evolution of castration-resistant neuroendocrine prostate cancer. Nat. Med. 22, 298–305 (2016).

54. Cancer Genome Atlas Research Network. Comprehensive molecular characterization of gastric adenocarcinoma. Nature 513, 202–209 (2014).

55. Subramanian, A. et al. Gene set enrichment analysis: a knowledge-based approach for interpreting genome-wide expression profiles. Proc. Natl. Acad. Sci. U. S. A. 102, 15545–15550 (2005).

