## Extended Data and Tables for "AGO1x prevents dsRNA-induced interferon signaling to promote breast cancer cell proliferation"

### *Extended Data Figures*

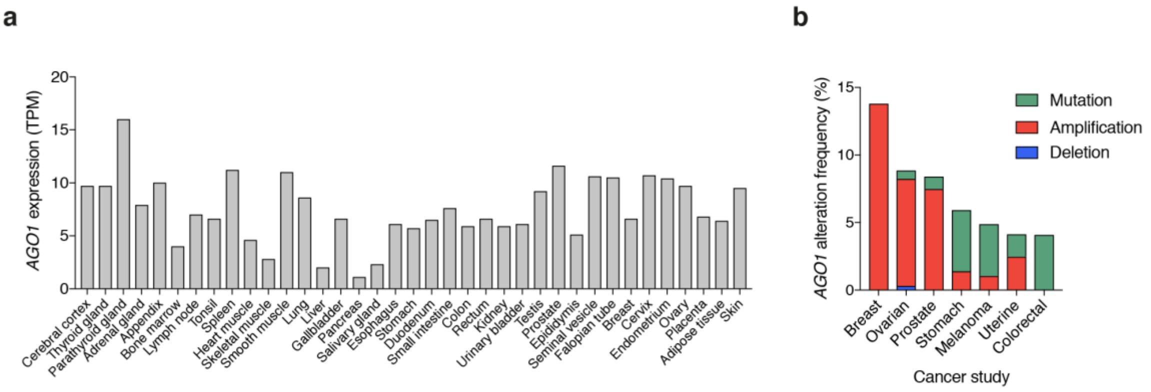


**Figure S1 | Expression profile of *AGO1* in normal tissues and genomic alterations of the locus in cancers.**

**a**, Expression level of *AGO1* transcript in tissues represented in the human protein atlas [14]. The height of the bars represents the mean expression across multiple samples (n=1-13, depending on the tissue). **b**, Primary tumor-derived tissues with highest frequency of alterations at the *AGO1* locus, retrieved (on February, 2018) from the cBio cancer genomics portal [15]: breast cancer xenografts [51], and various cancers: ovarian [52], prostate [53], stomach [54], melanoma (TCGA, provisional), uterine (TCGA, provisional) and colorectal (TCGA, provisional). Different types of genomic alterations are represented by different colors.

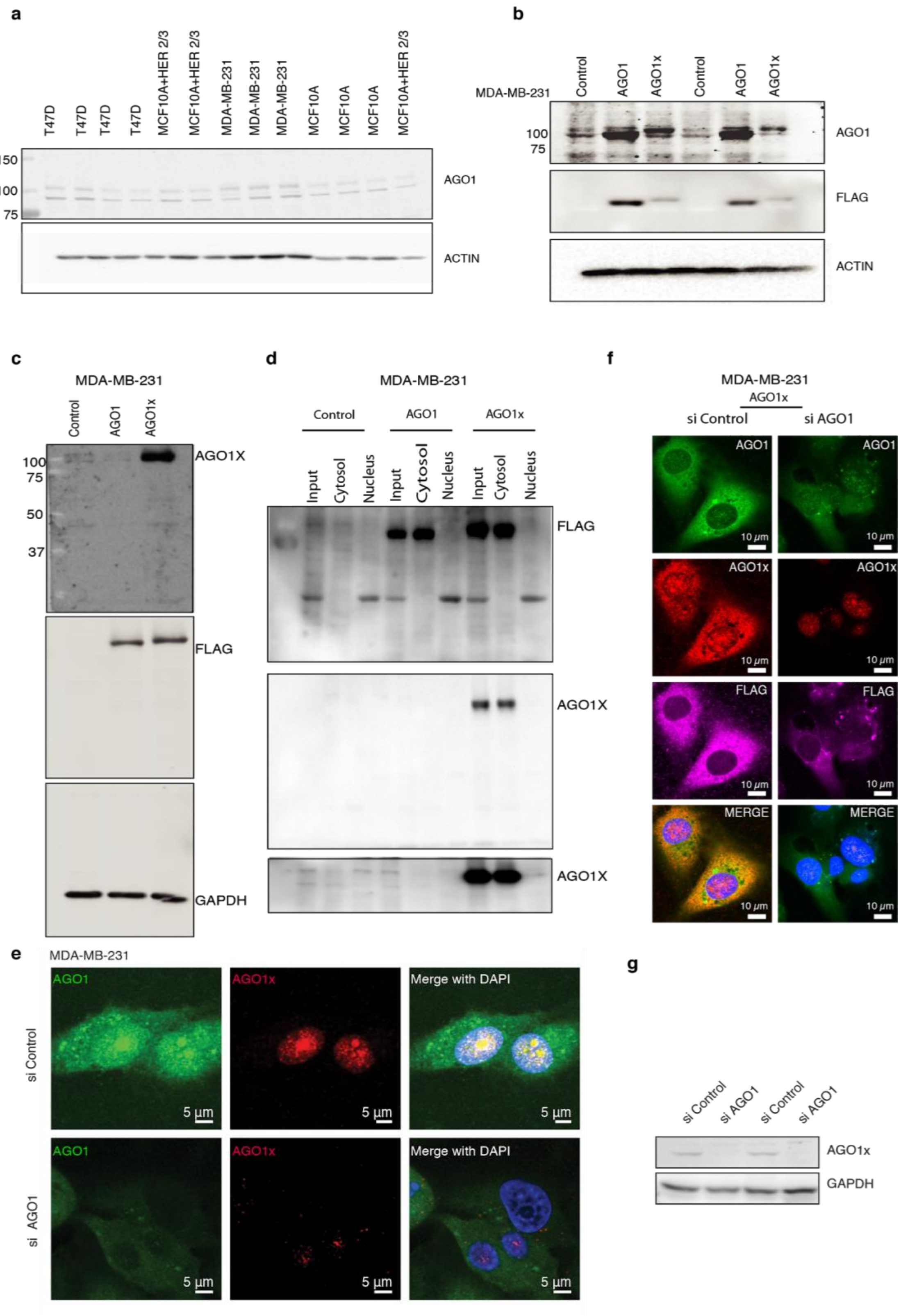
a
T47D
MCF10A+HER 2/3
MDA-MB-231
MCF10A
150
100
75
AGO1
ACTIN
b
MDA-MB-231
Control
AGO1
AGO1x
100
75
AGO1
FLAG
ACTIN
c
MDA-MB-231
Control
AGO1
AGO1x
100
75
50
37
AGO1X
FLAG
GAPDH
d
MDA-MB-231
Control
AGO1
AGO1x
Input
Cytosol
Nucleus
FLAG
AGO1X
AGO1X
f
MDA-MB-231
AGO1x
si Control
si AGO1
AGO1
AGO1x
FLAG
MERGE
10 μm
e
MDA-MB-231
si Control
si AGO1
AGO1
AGO1x
Merge with DAPI
5 μm
g
si Control
si AGO1
AGO1x
GAPDH

**Figure S2 | AGO1x antibody targets specifically its cognate protein and not the canonical AGO1**

**a,** Western blot analysis of multiple cell lines demonstrates a characteristic second band, at a higher MW, in addition to the canonical AGO1 protein band. **b,** Higher MW band observed in (**a**) corresponds to AGO1x protein. Representative western blot shows that the higher MW band intensity is sensitive to the ectopic overexpression (using the pIRES-Neo vector) of FLAG-tagged AGO1x but not of FLAG-tagged AGO1. Expression of the corresponding isoforms is confirmed with a blot for FLAG expression. Overexpression constructs used are indicated with labels above the blots. Protein ladders confirm that the higher MW band corresponds to a molecular weight of approx. 100 kDa. **c, d,** Western blot with AGO1x antibody shows its specificity for the AGO1x isoform stably expressed from pCDH-FLAG-tagged plasmids. **e-g,** Representative images of MDA-MB-231 stained with AGO1 (green) and AGO1x (red) antibodies at endogenous expression levels or under knock-down conditions with an siRNA pool targeting the transcript encoding both isoforms (**e**) or in systems of either AGO1 or AGO1x over-expression (**f**). DAPI was used to mark the nucleus (blue). (**g**) The levels of AGO1 and AGO1x proteins in cells pretreated with either siAGO1 or siControl were compared to the confocal imaging (left) by Western blot analysis.

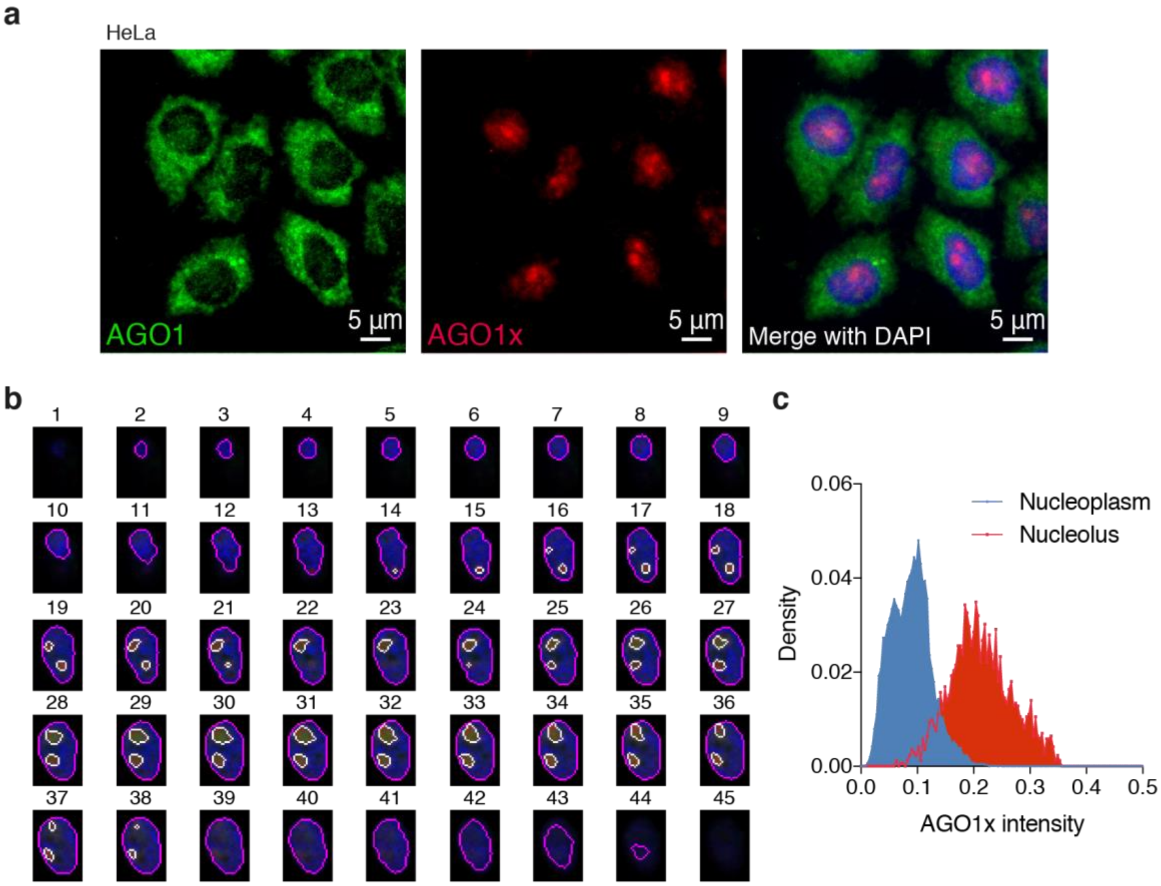


**Figure S3 | AGO1x localizes to the nucleus, in the vicinity of nucleoli.**
**a**, Immunofluorescence imaging of HeLa cells co-stained with AGO1 (green) and AGO1x (red) antibody. DAPI was used to mark the nucleus (blue). Images were documented with a Nikon Ti-E inverted microscope, cells were visualized with a CFI Plan Apochromat DM 60x lambda oil (NA 1.4) objective, and images were captured with a Hamatsu Orca-Flash 4.0 CMOS camera. **b**, Representative stacks of a MDA-MB-231 cell nucleus. The number of each z-stack layers is shown on the top of the image. DAPI, Nucleolin and AGO1x signals correspond to blue, red and green components of images, respectively. Detected outlines of the nucleus and nucleoli are depicted in magenta and white color, respectively. **c,** Histogram of AGO1x intensity in the areas corresponding to nucleoplasm and nucleoli from (**b**).

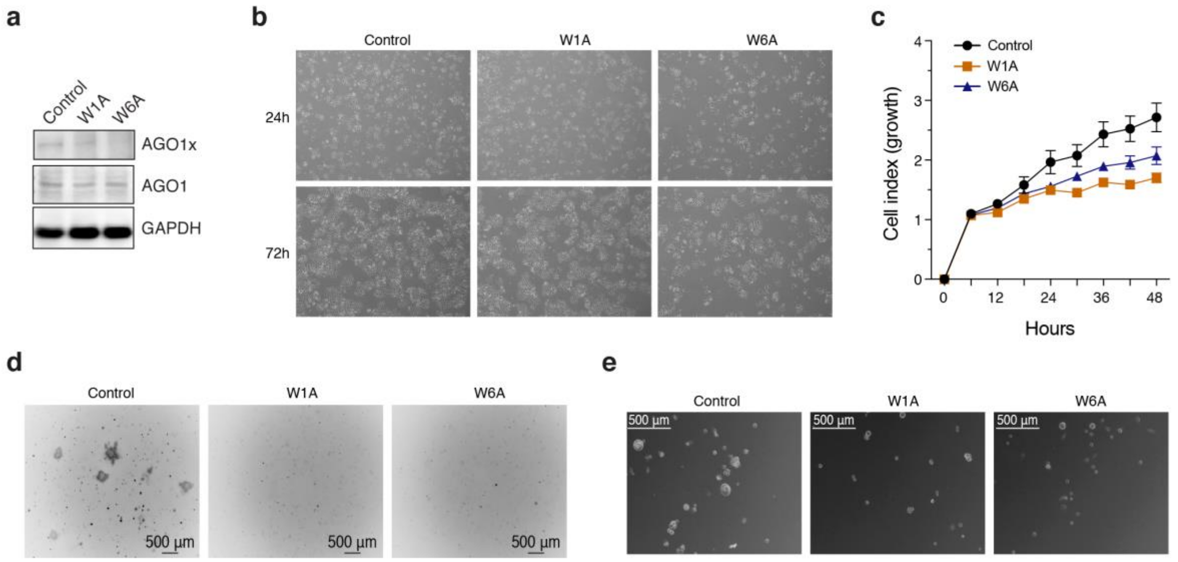


**Figure S4 | AGO1x deletion impairs cell growth.**

**a**, Western blot analysis of HeLa W1A and W6A mutant and control cell lines, using AGO1 and AGO1x antibodies. GAPDH served as an endogenous control for loading. **b**, Phase contrast representative images depict growth patterns of control and mutant HeLa cell lines at 24 hr and 72 hr after seeding equal numbers of each cell type in wells of a six well plate. **c**, Impedance-derived mean (+/- s.d.) cell indices at the indicated time points after seeding equal numbers of HeLa control (n=6), W1A (n=5) and W6A (n=5) cells. From 24 hours on, there is a statistical significant difference between control and the two mutants ($P < 0.005$, two tailed t-test). **d**, Representative images of colony formation assays performed with the W1A and W6A and control cell lines. Images were captured with an inverted microscope (ZEISS Axio Vert.A1 equipped with an AxioCam MRc camera). **e**, Representative images of sphere formation assays performed with MDA-MB-231 W1A and W6A mutant and control cell lines. Images were captured with an inverted phase contrast microscope (Leica) and were inverted for improved contrast.

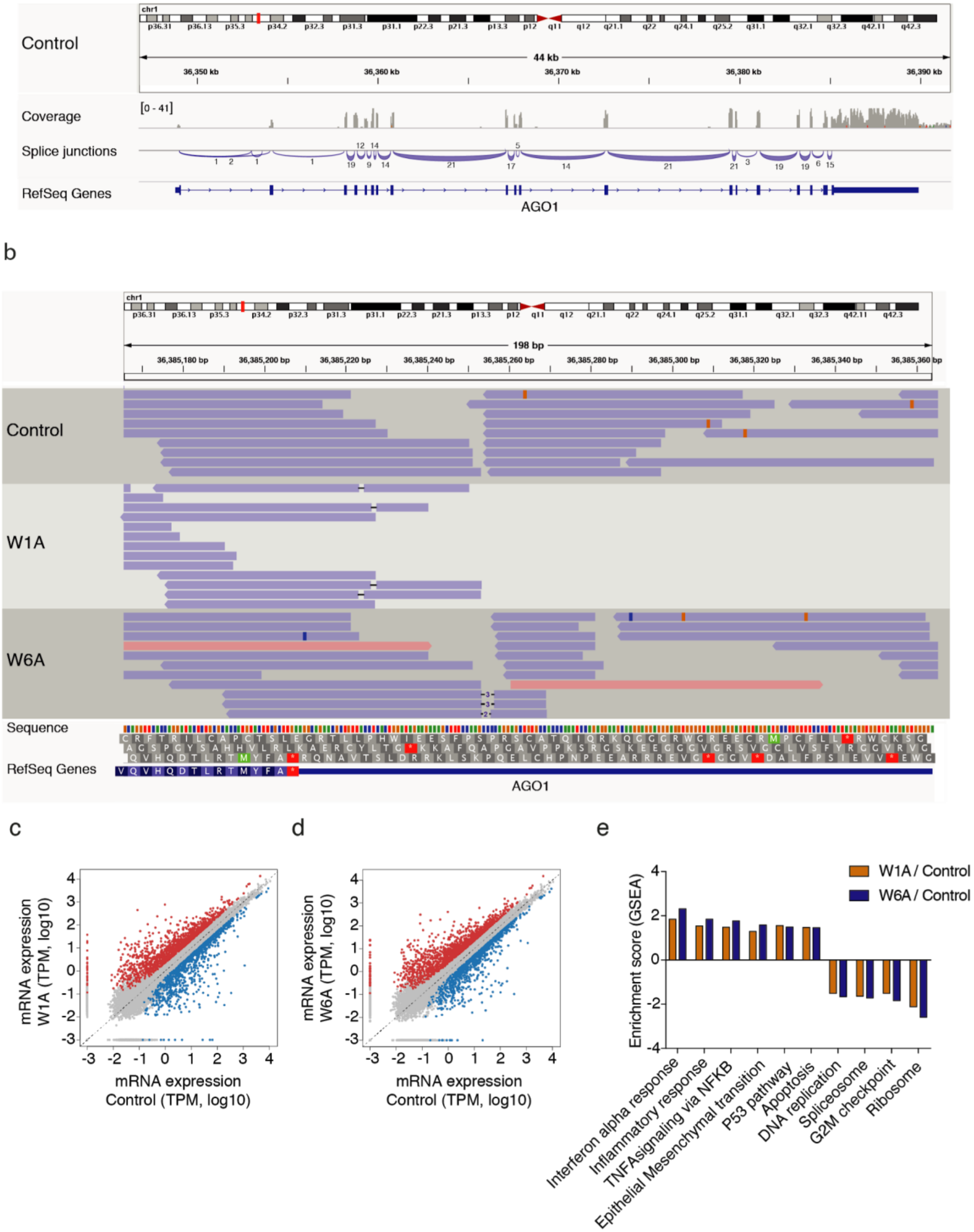

Control
chr1
44 kb
36,350 kb
36,360 kb
36,370 kb
36,380 kb
36,390 kb
Coverage
[0 - 41]
Splice junctions
RefSeq Genes
AGO1
b
198 bp
Control
W1A
W6A
Sequence
RefSeq Genes
AGO1
c
mRNA expression W1A (TPM, log10)
mRNA expression Control (TPM, log10)
d
mRNA expression W6A (TPM, log10)
mRNA expression Control (TPM, log10)
e
Enrichment score (GSEA)
W1A / Control
W6A / Control
Interferon alpha response
Inflammatory response
TNFAsignaling via NFKB
Epithelial Mesenchymal transition
P53 pathway
Apoptosis
DNA replication
Spliceosome
G2M checkpoint
Ribosome

**Figure S5 | Cellular pathways responding to AGO1x deletion.**

**a,** Read coverage profile and splice junction graph for the control cell line. The number of spliced reads that were sequenced is indicated for each splice junction. **b,** Alignment of RNA-seq reads from control and mutant cell lines samples to the region of TR that was targeted by the sgRNAs. Deletions observed in the RNA-seq from mutant lines are shown by dashes. **c**, **d**, Scatter plots of mRNA expression levels (log10 transcript counts per million, TPM) in W1A mutant (**c**) or W6A mutant cell lines (**d**) compared to control. Shown are mean expression values for each transcript (n=3). mRNAs that are significantly upregulated or downregulated (|fold-change| > 2-fold and FDR < 0.01) in the mutant lines are shown in red and blue, respectively. **e**, Normalized enrichment score (ES) derived from gene set enrichment analysis [55] comparing the expression of genes in the two mutant cell lines with that in the control cell line. For all the pathways depicted, $P < 0.05$. P-values were calculated by comparing the empirical ES of a gene set relative to a null distribution of ESs derived from permuting the gene set, and then adjusted for multiple hypotheses testing.

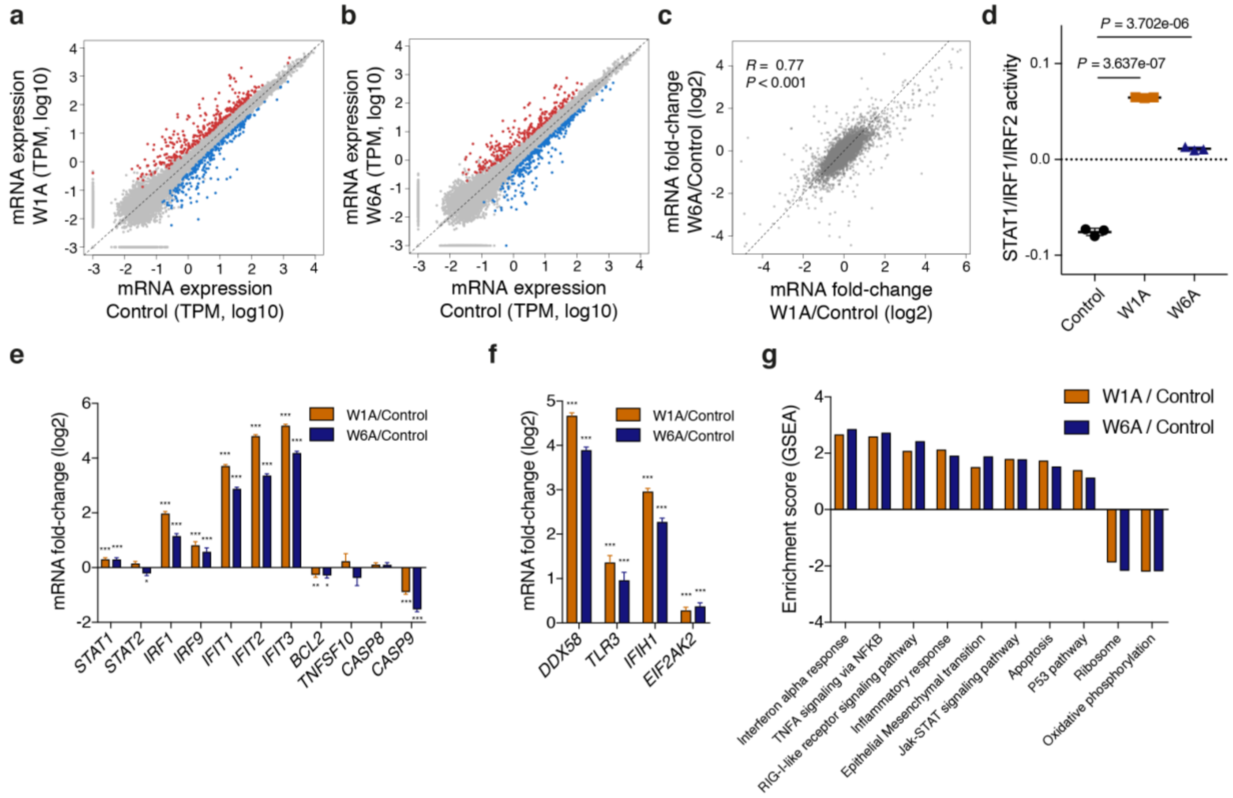

**Figure S6 | Deletion of AGO1x in HeLa cells activates the interferon response.**

**a**, **b**, Scatter plots of mRNA expression levels (log10 transcript counts per million, TPM) in W1A (**a**) and W6A mutant cell lines (**b**) compared to control. Shown are mean expression values computed for each transcript (n=3). mRNAs that are significantly upregulated or downregulated (|fold-change| > 2-fold and FDR < 0.01) in the mutant cell lines are shown in red and blue, respectively. **c**, mRNA fold-changes (log2) in the two mutant cell lines compared to control. Shown is also the Pearson correlation coefficient and respective P-value. Dashed line indicates equal fold-change in the two mutant lines. **d**, Mean (+/- s.d.) activity of STAT1/IRF1/IRF2 transcription factor motifs estimated by ISMARA [19] (n=3). Shown is also the P-value in an unpaired two-tailed t-test. **e**, **f**, Mean (+/- s.e.m.) mRNA expression fold-changes (log2) of several genes involved in the interferon alpha response and apoptosis (**e**) and in dsRNA sensing (**f**) in the two mutant cell lines relative to control (n=3). Multiple testing corrected P-values for fold-changes with respect to control are depicted above each bar (* $P < 0.05$, ** $P < 0.01$, *** $P < 0.001$). **g**, Normalized enrichment score (ES) derived from gene set enrichment analysis comparing the expression of genes in the two mutant cell lines as compared to the control cell line. For all the pathways depicted, $P < 0.05$. P-values were calculated by comparing the empirical ES of a gene set relative to a null distribution of ESs derived from permuting the gene set, and then adjusted for multiple hypotheses testing.

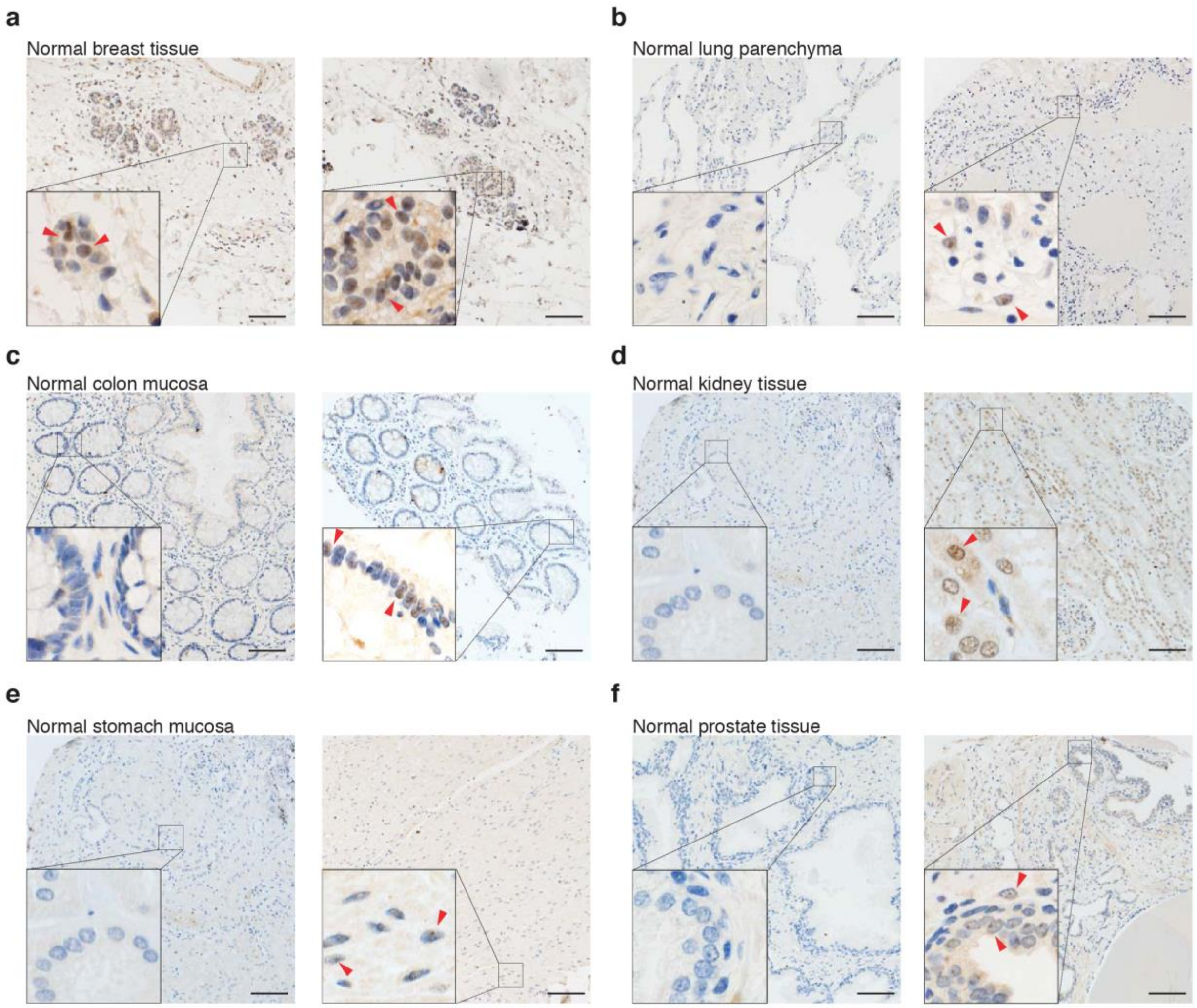


**Figure S7 | Immunohistochemical analysis of AGO1x expression in non-tumoral tissues**.
**a-f,** Representative micrographs of AGO1x expression in normal **a,** breast tissue; **b,** lung parenchyma; **c,** colon mucosa; **d,** kidney tissue; **e,** stomach mucosa, **f,** prostate tissue. Two representative non-tumoral tissue samples are shown for each organ. Insets show different degree of positivity at high magnification (red arrows) in the six different non-tumoral tissues. Scale bars, 100 μm.

*Extended Data Tables*

**Table S1 | Plasmids and oligonucleotides used in this study**.

| Name | Sequence Info | Company |
|---|---|---|
| si AGO1 | - | siTools Biotech #26523 |
| sgRNA W6A | AAGAAAGCTTTCCAAGCCCC | Microsynth AG |
| sgRNA W1A | GCAGAACGCTGTTACCTCAC | Microsynth AG |
| pSpCas9n(BB)-2A-Puro | - | Addgene # 62988 |
| pSpCas9(BB)-2A-GFP | - | Addgene # 48138 |
| pIRES-Neo-Flag/HA-AGO1 | - | Addgene # 6798 |
| pIRES-Neo-Flag/HA-AGO1x | Modified from Addgene #6798,<br>Additional nucleotides appended to the stop codon in #6798: aggcagaacgctgttacctcactggatagaagaaagctttccaagccccaggagctgtgccacccaaatccagaggaagcaaggaggagggaggtggggtag. The stop codon of #6798 modified to tcc. | - |
| pCDH-AGO1 | AGO1 sequence from pIRES-Neo-Flag/HA-AGO1 was cloned into the pCDH expression vector | System Biosciences #CD527A-1 |
| pCDH-AGO1x | AGO1x sequence from pIRES-Neo-Flag/HA-AGO1x was cloned into the pCDH expression vector | - |
| ON-TARGETplus Human PNPT1 (87178) siRNA | - | Horizon Discovery |
| FDFT1 Fwd | GACACGTGGGCGACTTATTG | Microsynth |
| FDFT1 Rev | AGCGAGTCCTGGTCCATCTT | Microsynth |
| SCD Fwd | TCCAGAGGAGGTACTACAAACCT | Microsynth |
| SCD Rev | CCGGGGGCTAATGTTCTTGT | Microsynth |
| EFHD2 Fwd | GATGTTCAAGCAGTATGATGCC | Microsynth |
| EFHD2 Rev | CTTGCGGAAGATCAGGAGGAA | Microsynth |
| SEMA6B Fwd | ACCGTTGTCTTCCTGGGTTC | Microsynth |
| SEMA6B Rev | GCCGATACAGTTCTTCATACACC | Microsynth |
| HAS2 Fwd | GCTGAACAAGATGCATTGTGAG | Microsynth |
| HAS2 Rev | ATAGGCAGCGATGCAAAGGG | Microsynth |
| 5.8 S rRNA Fwd | CTTAGCGGTGGATCACTCGG | Microsynth |

| | | |
|---|---|---|
| 5.8 S rRNA Rev | AGTGCGTTCGAAGTGTCGAT | Microsynth |
| 28S rRNA Fwd | AGGTAGCCAAATGCCTCGTC | Microsynth |
| 28S rRNA Rev | TTCACCGTGCCAGACTAGAG | Microsynth |
| 18S rRNA Fwd | TGACTCTAGATAACCTCGGG | Microsynth |
| 18S rRNA Rev | GACTCATTCCAATTACAGGG | Microsynth |
| 45S pre-ribosomal RNA Fwd | GTGAAACCTTCCGACCCCTC | Microsynth |
| 45S pre-ribosomal RNA Rev | TACGAGGTCGATTTGGCGAG | Microsynth |
| NUDT21 Fwd | TGTACATGAGCACCGGCTAC | Microsynth |
| NUDT21 Rev | CCTGACGACCCAGTATCTCTG | Microsynth |
| CPSF6 Fwd | TGAGTCCAAGTCTTATGGTTCTGG | Microsynth |
| CPSF6 Rev | CCTCTTCCTTCAGCTTCTAACGA | Microsynth |
| GAPDH Fwd | AATCCCATCACCATCTTCCA | Microsynth |
| GAPDH Rev | TGGACTCCACGACGTACTCA | Microsynth |

**Table S2 | Antibodies used in this study**.

| **Name** | **Company** |
|---|---|
| α-Tubulin | Calbiochem, # CP -06 |
| Nucleolin | Thermo Scientific, # 396400 |
| ERP72 | BD Biosciences, # 610970 |
| Lsm4 | Sigma, # GW22314F |
| SC35 antibody | Abcam, # ab11826 |
| AGO1x | Lucerna-Chem, # RBP 1510 |
| AGO1 | Novus Biologicals, # NB100-2817 |
| p54 | BD Biosciences, # 611278 |
| GAPDH | Sigma, # G9295 |
| hnRNP C1/C2 | Santa Cruz, # SC10037 |
| Ki-67 | Dako, # IR626 |
| J2 | Scicons # 10010200 |
| PNPT1 AB (4C11) | Novus #NBP2-43725 |
| DHX 9 | Bethyl #A300-855A |
| ANTI-FLAG® M2 | Sigma # F3165-.2MG |
| Actin | Santa Cruz #SC1615 |
